# Comparing Symbiodiniaceae diversity across unicellular hosts, multicellular hosts, and environmental reservoirs

**DOI:** 10.64898/2026.04.21.719777

**Authors:** Vera Emelianenko, Maria E. A. Santos, Gayun Kim, Filip Husnik

**Affiliations:** Okinawa Institute of Science and Technology; Evolution, Cell Biology, and Symbiosis Unit 1919-1 Tancha, Onna-son, Okinawa 904-0495, Japan; Hawai‘i Institute of Marine Biology, University of Hawai‘i at Mānoa, Hawai‘i, U.S.A

## Abstract

Symbiodiniaceae dinoflagellates are the primary photosymbionts in reef ecosystems, crucial for reef productivity. Although they are widely recognized as symbionts of animals such as corals and clams, they can occupy a broad range of reef niches, including water, sediment, and macroalgae. Understanding their ecology is typically hampered by their horizontal acquisition. Combining evidence from multiple sample types collected at the same location has the potential to address this issue, but such analyses are surprisingly rare. Here, we analysed Symbiodiniaceae communities across 74 environmental and host samples in one reef flat in Okinawa, Japan. We detected ten Symbiodiniaceae genera or genus-level clades using the ITS2 marker metabarcoding, including Clade J, previously known only from Okinotori Island, Japan. *Cladocopium, Symbiodinium*, and *Durusdinium* dominated multicellular hosts (hexacorals and *Tridacna*). In contrast, foraminiferal hosts were dominated by *Cladocopium* or genus-specific *Freudenthalidium, Fugacium*, and *Miliolidium*. Symbiont communities were mostly specific to the host genera. Water samples, with higher proportions of *Durusdinium* and free-living *Symbiodinium*, were distinct from macroalgae and sediment samples. The latter did not differ significantly from each other and contained *Freudenthalidium, Fugacium, Miliolidium*, Clade I, and Clade J. Only three ITS2 variants were shared across all sample categories, but many variants were unique to hosts or habitats. We highlight that both unicellular and multicellular hosts harbor specific endosymbiont types, with lower diversity than in the surrounding environment. Our results imply that host diversity, availability, and environmental context jointly structure photosymbiont communities at fine spatial scales within coral reef ecosystems.

## Introduction

Symbiodiniaceae (Suessiales, Dinoflagellata) are the most common photosymbionts of eukaryotes, forming symbioses with a wide range of both multicellular (corals, jellyfish, and giant clams) and unicellular hosts (foraminiferans and ciliates) [1]. Originally, they were described as *Symbiodinium* with clades A-I, but the clades were elevated to the genus level [2] due to their high molecular and morphological diversity. Apart from mutualistic associations, Symbiodiniaceae can also be free-living or opportunistic along the symbiosis continuum [1]. While many studies describe Symbiodiniaceae associated with corals, Symbiodiniaceae diversity and ecology in unicellular hosts and free-living environments have received much less attention [3–5]. However, microhabitats such as sediment, water, and macroalgae serve as reservoirs for symbionts, which hosts acquire horizontally during development or following bleaching events. Characterizing the free-living lineages of Symbiodiniaceae related to the host-associated ones can not only help us understand the adaptations to symbiosis with different hosts, but also provide us with comparative experimental models. Here, we compared Symbiodiniaceae communities associated with animals (clams and hexacorals), Foraminifera, and the surrounding environmental samples, to discuss symbiont specificity and potential acquisition reservoirs within the same reef flat.

## Results

In total, our analyses of the Symbiodiniaceae communities in six animal genera, two foraminiferal genera, and three types of environmental samples detected 111 Symbiodiniaceae amplicon sequence variants (ASVs) of the internal transcribed spacer 2 (ITS2) of the rDNA region, spanning seven Symbiodiniaceae genera and three genus-level clades (Fig. 1). Corals and clams were associated with *Symbiodinium, Cladocopium*, and *Durusdinium*, while Foraminifera were dominated by *Freudenthalidium, Fugacium, Miliolidium*, or *Cladocopium*. All the genera and genus-level clades we detected were also present in environmental samples, although overall Symbiodiniaceae counts were low and thus less reliable. Clades I–J were consistently detected in sediment and macroalgae. In contrast, water showed a high proportion of *Durusdinium* together with *Cladocopium* and *Symbiodinium*, while having no or few sequences of other genera.

**Figure 1.**
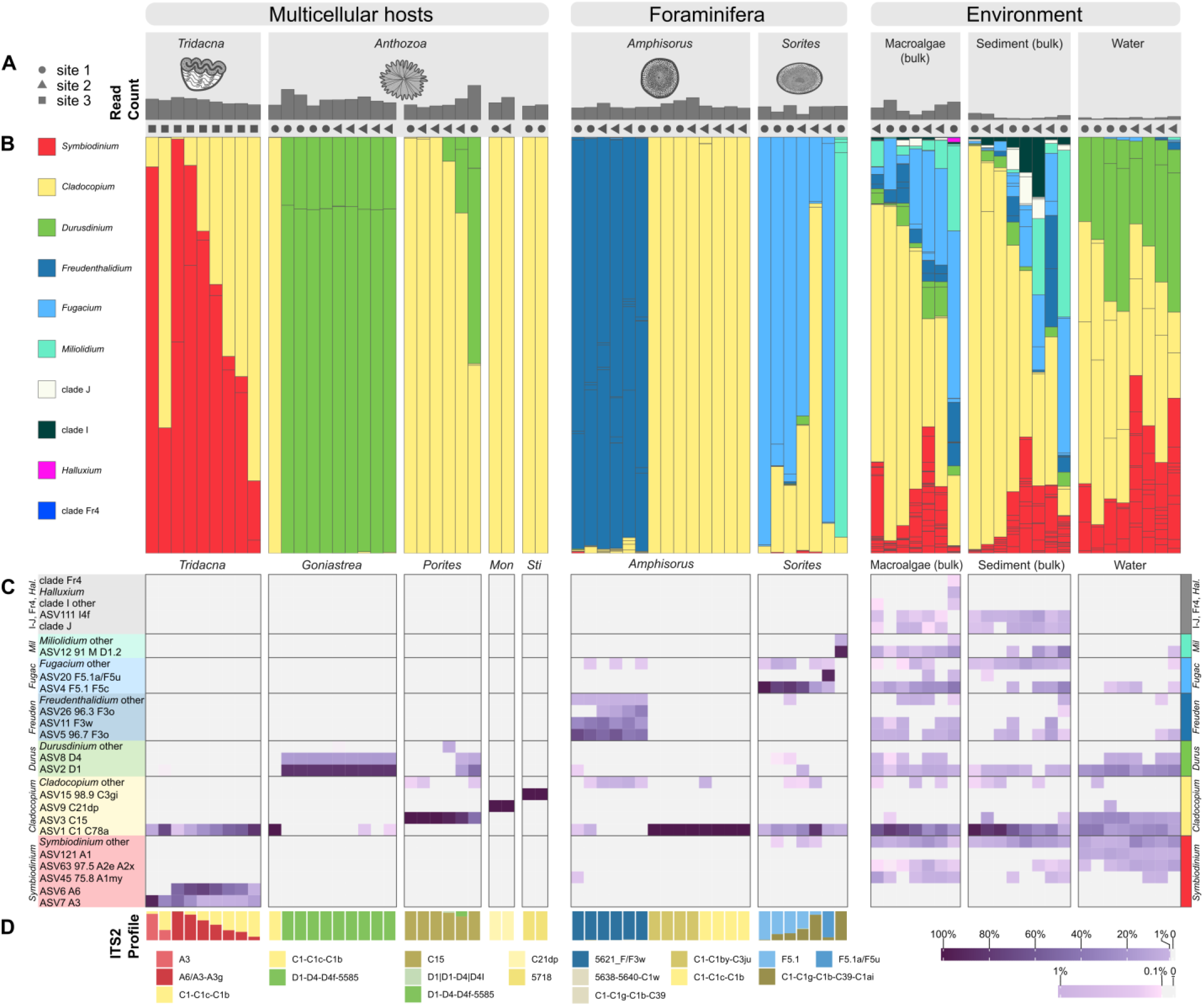
Symbiodiniaceae ITS2 composition across hosts and environments. A – Total count of reads mapped to Symbiodiniaceae ASVs (see Table S2 and Fig. S1 for details). B – Bar chart displaying distribution of Symbiodiniaceae ITS2 sequence variants assigned to genera (labeled by colours). The abundance of corresponding genera may differ due to ITS2 copy number variation. Different geometric shapes (circles, triangles, and squares) denote the sampling sites. *Montipora* samples are labeled by “*Mon*” and *Stichodactyla* by “*Sti*”. Thin grey lines separate individual ASVs within a bar. C – Heatmap displaying relative abundance of ASVs. The variants that constitute >10% in at least one sample are shown separately; the remaining ones are collapsed by genus. Only abundances >0.1% are shown. The labels show ASV identifier followed by the highest identity match in the SymPortal named database, with identity shown if <100%. D – ITS2 profiles of Symbiodiniaceae for host samples, based on SymPortal analysis. ITS2 profile is a combination of defining intragenomic variants that define a taxonomic unit according to SymPortal. Each column in A–D represents the same sample. One of the macroalgal samples consisting only of C1 *Cladocopium* was removed as an outlier (see Figs. S1, S2). All *Tridacna* samples belong to the species *T. maxima*, except Tridacna1 (second column), which belongs to *T. noea*. See Fig. S9 for details of the identification of Foraminifera.

Symbiodiniaceae community composition was influenced by sample type (Fig. 2A), which explained about 80% of the variation among samples collected on the reef flat from sites 1 and 2 (PERMANOVA: *R*^2^ = 0.805, *F* = 28.43, *p* < 0.001; PERMDISP: p > 0.05; consistent when analyzed using SymPortal [6] Defining Intragenomic Variants [DIVs]). In contrast, the sampling site had no significant effect (R^2^ = 0.012, *F* = 0.76, *p* = 0.511). Symbiodiniaceae community composition of water samples differed from both algal and sediment samples (R^2^ = 0.708, *F* = 31.51, False Discovery Rate (FDR)-adjusted *p* = 0.002, and R^2^ = 0.640, *F* = 24.86, FDR-adjusted *p* = 0.002, respectively), while algal and sediment sample types had no significant difference (Table S3). Although multicellular hosts showed a broad spread along the ordination axes, their separation was primarily driven by differences in the proportions of unique *Symbiodinium* types in clams and *Durusdinium* in anthozoans (Figs. 2B, S5). In contrast, *Cladocopium* had comparatively little effect on the sample type separation, as it was present across all sample types and was represented by phylogenetically close sequences.

**Figure 2.**
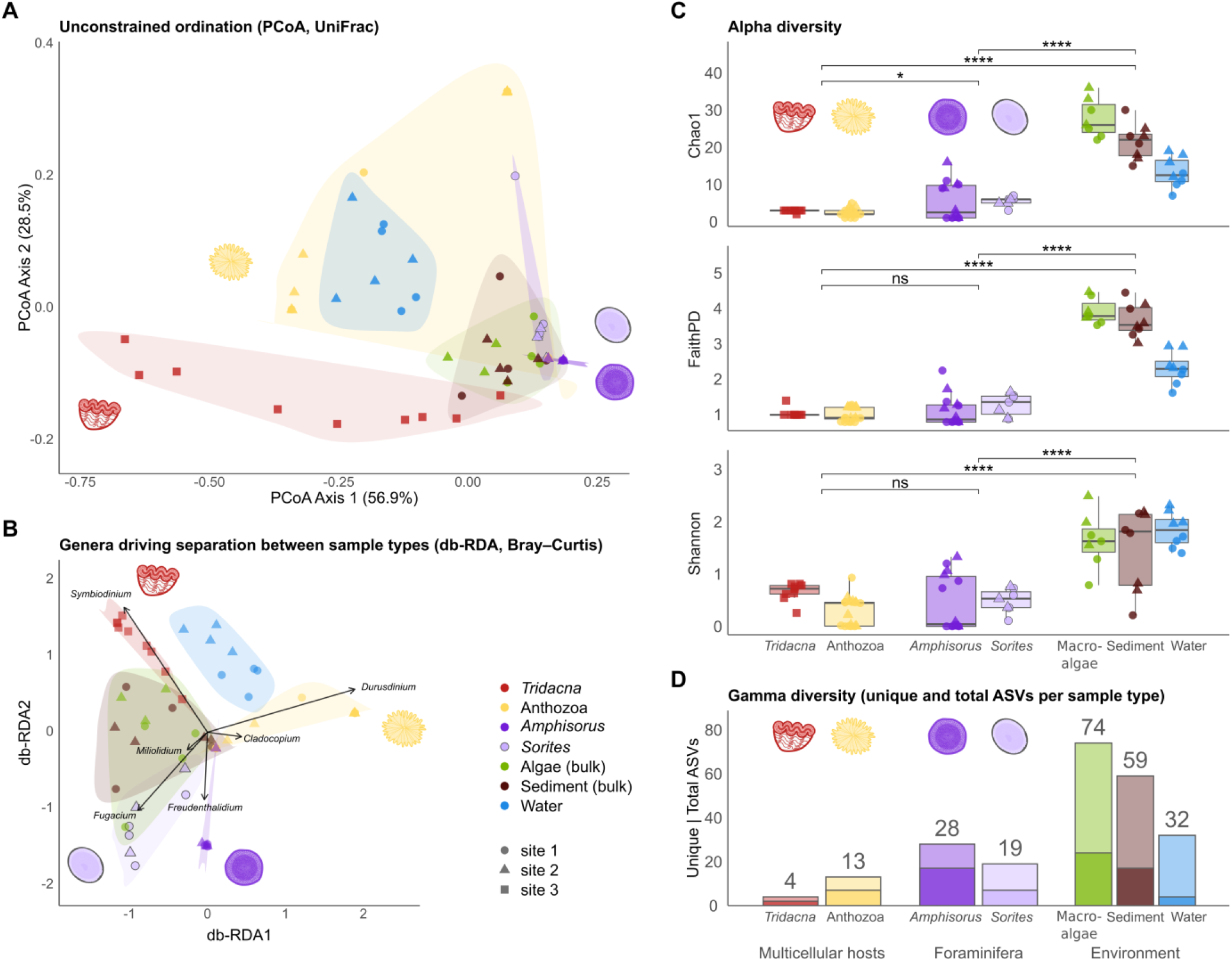
Diversity metrics and comparison between the habitats. A – Principal coordinates analysis (PCoA) based on UniFrac distances (considers phylogeny and relative abundance), with sample type labeled by colour and sampling sites labeled by shape. B – Distance-based redundancy analysis (db-RDA) based on Bray–Curtis dissimilarities, aggregated at the genus level, constrained by sample type; black arrows indicate the six genera most strongly associated with the separation among sample types. Overlapping purple points in the middle of A and B represent multiple samples with *Cladocopium* C1 and closely related ITS2 variants as their only symbiont. C – Diversity indices based on ITS2 sequence data of the unfiltered counts (effects of filtering and rarefaction are shown in Fig. S4). From top to bottom: Chao1, Faith’s phylogenetic diversity (PD), and Shannon diversity index. D – Total number of ASVs (values shown above bars) with unique ASVs (lines in the middle) per sample type. “Anthozoa” represents multiple genera (*Goniastrea, Montipora, Porites, Stichodactyla*). Phylogenetic trees used to calculate distances were approximated separately using FastTree on unfiltered (Faith’s PD) and filtered (PCoA) data. Comparisons with SymPortal data are shown in Fig. S3.

Diversity metrics consistently showed higher values for the environmental samples compared to the hosts. Alpha diversity metrics (Figs. 2C, S5A) were significantly higher in environmental samples regardless of pipeline, filtering, and rarefaction steps (Wilcoxon test FDR-adjusted p < 0.05: Fig. S4, Table S4). The total unique ASVs and DIVs per type showed the same pattern (Figs. 2D, S6B). In contrast, no or little difference in per-sample alpha diversity measures was detected between multicellular hosts and Foraminifera (all Wilcoxon test FDR-adjusted p > 0.025, most comparisons > 0.05; Table S4). However, Foraminifera demonstrated a higher number of unique ASV per sample type than multicellular hosts. Richness accumulation curves based on ASVs and DIVs suggested that not all the potential diversity in environmental samples was sampled (Fig. S6).

Shared and unique ASVs were observed across sample types (Figs. 2D, S7), with each sample type having unique ASVs. Only *Cladocopium* C1 and *Durusdinium* D1-D4 were shared between all the groups: multicellular hosts, Foraminifera, and environmental samples. Unsurprisingly, Symbiodiniaceae communities overlapped between Foraminifera and bulk macroalgal and sediment samples (Fig. S7). Macroalgae and sediment are potential symbiont reservoirs for Foraminifera [7]. Still, this overlap can be partially explained by small Foraminifera in those samples. However, Symbiodiniaceae composition of both foraminiferal genera was distinct from macroalgae and sediment (Table S4). Many ITS2 variants were detected only in the environment, including Clades I –J, Fr4, *Halluxium, Fugacium* F5.2, and 31 *Symbiodinium* ASVs, with some variants specific to water samples (A1). Unlike variants of *Durusdinium*, environmental *Symbiodinium* variants did not overlap with host-associated ones.

## Discussion

Horizontal acquisition of symbionts from the environment is, perhaps counterintuitively, the rule rather than the exception in the marine environments. Due to the transmission mode, major model systems such as the squid-*Vibrio* bioluminescent symbiosis [8], chemosynthetic symbioses of hydrothermal vent mussels [9], and coral-dinoflagellate photosymbioses [10,11] rely on the presence of the potential symbionts in the environment. However, the distribution and diversity of the free-living lineages in the environment are understudied, and the host-symbiont specificity is often context-dependent. Additionally, some symbionts, including Symbiodiniaceae, exhibit both vertical and horizontal transmission depending on the host. In our data, host-symbiont pairs were largely consistent with previously published data from the Indo-Pacific region [12–18]. Two genera of corals analysed in this study have vertical transmission of symbionts, and accordingly host-specific symbionts: the C15 variant in the Indo-Pacific *Porites* [2,13] and the C21-related variant in *Montipora* [13,14]. Similarly, giant sea anemones are known to have vertical symbiont transmission and specific symbionts not shared with scleractinian corals, which explains why we did not detect *Stichodactyla*’s symbionts in other hosts or environments and why their symbionts’ ITS2 sequences from C3 lineage were not present in the database. Type C1, present in all sample types, was previously recorded from Indo-Pacific *Tridacna maxima* [14], *Goniastrea* [14], and Foraminifera [15]. Although it appears as a host-generalist, the C1 variant is in fact characteristic of several *Cladocopium* species [19]. *Symbiodinium* A3–A6 variants in giant clams are consistent with *S. tridacnidorum*, and D1–D4 variants are characteristic of *D. trenchii* [20], common in Indo-Pacific corals and known for thermal tolerance. Unexpectedly, we found *Miliolidium* in a *Sorites* sample (Fig. S9), although this symbiont genus is primarily associated with *Marginopora* [21].

The higher diversity of environmental samples compared to host-associated ones is consistent with previous observations [5]. Among several potentially free-living clades, Clade J was previously described only from the Okinotori Island [22]. In addition, multiple divergent *Symbiodinium* ITS2, putatively free-living, were also present in environmental samples [7,22,23]. Most sequencing data in environmental samples originated from non-Symbiodiniaceae ASVs (Fig. S1), explaining the low total Symbiodiniaceae counts (Fig. 1A). High proportions of ASVs assigned to the order Suessiales point to undescribed free-living Suessiales diversity in these microhabitats.

In contrast to previously published data [3], we found no separation between macroalgal and sediment samples, while water samples were distinct. We also did not detect variability between sampling sites, even though environmental Symbiodiniaceae communities demonstrate extensive small-scale variability [4,5]. Although we included several potential symbiont reservoirs, further research should also examine biological vectors, such as coralivorous fish feces [24].

## Conclusion

Our results show that all host types were associated with distinct Symbiodiniaceae, which were only partly shared with environmental samples. Environmental samples consistently demonstrated higher diversity of Symbiodiniaceae ASVs than host samples, with water Symbiodiniaceae communities distinct from those in macroalgae and sediment. Considering the community variability between studies, future work should focus on experimentally identifying the specific factors that shape the distribution of Symbiodiniaceae clades.

## Data availability

The raw sequenced data of Symbiodiniaceae ITS2 obtained in this study have been deposited in the NCBI Sequence Read Archive under the BioProject ID PRJNA1299766. GenBank accession numbers of the gene sequences used for the host identification are PV984354-PV984355, PV984519-PV988047, PX026258-PX026260, and PX092634-PX092652. Alignments and photos were uploaded to FigShare 10.6084/m9.figshare.29881454. Scripts used to analyse the data and count tables were uploaded to GitHub https://github.com/ECBSU/ECBSU_manuscripts_code/tree/main/Eme-lianenko_Comparing_Symbiodiniaceae_diversity-2026.

## Acknowledgements

We are grateful for the help and support provided by the Sequencing Section of Core Facilities at OIST, especially for the library preparation and Illumina sequencing, the OIST Scientific Computing & Data Analysis Section for providing high-performance computing resources and storage, Dr. Dewi Langlet for advice and feedback on foraminiferal symbionts, Kenta CF Kondo for the identification of the sea anemone, and Sachie Matsuoka for assistance with sampling logistics. F.H.’s group was partially supported by OIST institutional funding, the COI-NEXT program, and an HFSP Early Career Research Grant [https://doi.org/10.52044/HFSP.RGEC292024.pc.gr.194160].

## Author contributions

V.E., M.S., and F.H. conceptualized and designed the research project. V.E. and M.S. collected samples. V.E. performed sample processing, DNA extractions, and data analysis. V.E. and G.K. identified the hosts with molecular markers. V.E. prepared the first draft of the manuscript. M.S. and F.H. provided feedback and revised the manuscript. F.H supervised the project.

## Materials and Methods

Our sampling included Foraminifera *Amphisorus* sp. and *Sorites* sp., corals *Goniastrea* sp., *Porites* sp., and *Montipora* sp., a sea anemone *Stichodactyla* sp., and environmental samples of water, sediment, and miscellaneous macroalgae at Ikei-jima reef flat in Okinawa, Japan (see Suppl. methods for details of host identification). The samples were collected by reef-walking from two sites, separated by ∼200 m, on 24 February 2023. We additionally analysed giant clams *Tridacna* (*T. noeae* and *T. maxima*) from the deeper part of the same reef, which we purchased from fishermen in the same month. After DNA extractions, the internal transcribed spacer 2 (ITS2) of the rDNA region, the marker most frequently used to assess Symbiodiniaceae lineages [20], was amplified with Symbiodiniaceae-specific primers ITS-DINO [25] and ITS2Rev2 [26]. We quantified abundances of amplicon sequence variants (ASVs) in all samples with the DADA2 pipeline [27] and LULU curation [28] and assigned genera using a database modified from [3] and SymPortal [6]. Additionally, we used SymPortal to assign Defining Intragenomic Variants (DIVs) to all samples and ITS2-type profiles to host samples.

## Supplementary methods

### Sample collection

Environmental and host samples, except for *Tridacna*, were collected on 24 February 2023 on Ikei Island (Okinawa, Japan) by reef walking. Two sites, separated by ∼200 m, were used for collection: site 1 (26.393975, 128.002729) and site 2 (26.392062, 128.002119). *Tridacna* were purchased from the Yonashiro Fisherman’s Association, who collected them on 2 February 2023, collection coordinates: 26.396019, 128.008487; 26.391502, 128.006234; 26.386543, 128.004903. *Tridacna* tissue was sampled from the mantle with sterilized scissors. Coral samples around 1 cm^3^ in volume, identified as *Goniastrea* sp. and *Porites* sp. in the field, were collected with a chisel and hammer (coral sampling permit number 4-15). Soritid foraminifera, *Amphisorus* sp. and *Sorites* sp., and sea anemone *Stichodactyla* sp. were picked from rocks upon returning to the laboratory with a fine paintbrush or forceps. Before picking, rocks were kept in closed aquaria with seawater for up to 1 week. Foraminifera were checked for pseudopodia to confirm they were alive, photographed, washed in filtered seawater three times, and kept at −20 °C until further processing. We collected sediment samples in 50 mL falcon tubes (*n* = 8 per site), miscellaneous macroalgal samples in plastic bags (*n* = 4 per site), and water samples in 5 L bottles (*n* = 4 per site). Water was gravity-filtered through the 42 μm nylon mesh to prevent large particles and zooplankton accumulation, then vacuum filtered through the 5 μm polytetrafluoroethylene (PTFE) Omnipore filter (Millipore). Filtration was performed on the same day as the collection. The filters were stored at −70 °C until DNA extraction. All other samples were stored at −20 °C until further processing.

### DNA extraction and library preparation

Macroalgal and sediment samples (50 mL of sediment or 50 g of macroalgae) were ground for 10 min using the Mortar Mill RM200. Total DNA was extracted from two ∼0.5 mL subsamples of the ground mass, which were processed separately until data analysis. DNA from host samples, sediment samples, and macroalgal samples was extracted using the DNeasy PowerSoil Pro Kit (QIAGEN) according to the manufacturer’s protocol, with incubation at 65 °C for 10 min for macroalgal, sediment, and foraminiferal samples. Foraminiferal shells were crushed with UV-sterilized stainless-steel nails before DNA extraction. DNA from the water samples was extracted using the DNeasy PowerWater Kit (QIAGEN). Extraction tubes without adding tissue or ground material were used as negative controls for DNA extraction.

DNA concentration was quantified using the Invitrogen Qubit DNA HS kit (Thermo Fisher Scientific), and samples with DNA concentrations lower than 0.1 ng/μL were not used further. Therefore, for some sample categories, the number of samples processed was lower than initially aimed (for example, only 3 *Sorites* samples were submitted for site 2). For each category, 4–8 samples were submitted to the Okinawa Institute of Science and Technology Sequencing Section for library preparation and sequencing. ITS2 region of the rRNA gene was amplified with Symbiodiniaceae-specific primers fused with overhangs: ITS-DINO 5’-TCGTCGGCAGCGTCAGATGTG-TATAAGAGACAG-GTGAATTGCAGAACTCCGTG-3’ [1] and ITS2Rev2 5’-GTCTCGTGGGCTCGGAGATGTGTATAAGAGACAG-CCTCCGCTTACTTA-TATGCTT-3’ [2] (dash indicates the border between the primer and the overhang sequence for index primers). PCR was performed in a 20 μL volume with 1 μL of each primer (5 μM), 3 μL of DNA normalized to 12.5 ng, and 12.5 μL of 2x KAPA HiFi HotStart ReadyMix, with the following conditions: initial denaturation at 98 °C for 30 s, followed by 30 cycles of 98 °C for 10 s, 52 °C for 30 s, and 72 °C for 30 s, with a final extension step of 5 min at 72 °C. Elution buffer (10 mM Tris-HCl, pH 8.0) was used as the negative control for the PCR reactions. PCR products were cleaned with AMPure XP beads (Beckman Coulter), and 5 μL was used in the second PCR reaction with 2 μM Nextera XT Index Primers and 2x KAPA HiFi HotStart ReadyMix. The second PCR products were cleaned with AMPure XP beads and sequenced on two lanes of Illumina MiSeq with 300 bp paired reads using a version 3 (600 cycle) reagent kit.

### Data analysis

The primary analysis — including BCL conversion, demultiplexing, and adapter trimming — was performed using bcl2fastq v2.20 at OIST Sequencing Section, with AdapterRead1 = CTGTCTCTTATACACATCT, AdapterRead2 not specified, and default adapter stringency (0.9). To trim primers from raw sequences and discard the sequences not containing both forward and reverse primers, reads with ambiguous bases were discarded, and cutadapt 4.2-1 [3] was used with 20% mismatch allowed in primer matching. Quality filtering was performed with the DADA2 v1.26.0 pipeline [4] using “filterAndTrim” allowing for a maximum of 2 expected errors in forward reads and 4 errors in reverse reads to retain otherwise low-quality reverse reads; only reads shorter than 50 bp long were removed to allow for a variable size of the ITS2 region; PhiX reads were removed with DADA2’s default filter. The number of reads per sample during different stages of quality filtering was assessed with seqkit 2.3.1 [5]. Amplicon Sequence Variants (ASVs) were assigned with DADA2. Data from three different sequencing runs were processed separately and merged using the “mergeSequenceTables” function after the ASV assignment step. The merged sequence table was used to remove chimeric reads. ASVs with a minimal length of 250 bp were selected using seqkit. The length cut-off was based on visual inspection of the length histogram and average size of Symbiodiniaceae ITS2 amplicon, but was specifically set liberally to keep as many ASVs as possible (compare to the length cut-off of ∼294–304 bp in [6]). Symbiodiniaceae composition in separate aliquots of macroalgae and sample sediment was assessed with a bar chart (Fig. S2), and then counts for the same sample were merged between different aliquots and different runs. Batch effects were not checked as the first two runs represented the same set of samples, with the third run contributing only two *Amphisorus* samples with the same Symbiodiniaceae composition as previously sequenced *Amphisorus*. Co-occurring ASVs with high identity were clustered with LULU 0.1.0 [7], with a minimum identity and a minimum relative co-occurrence 95%. Match list for LULU curation was created with BLASTn 2.12.0 with minimal query coverage of 80% and minimal identity of 95%, using the same set of ASV sequences as the database and the query. ASVs retained by LULU (“parents”, 922 retained and 512 discarded) were used in the downstream analysis together with the curated ASV table produced by LULU.

Taxonomy was assigned to curated ASVs with DADA2 1.36.0 “assign-Taxonomy” function with a minimum bootstrap of 50. To ensure the most precise genus assignment, the database was modified from the database from [8] by adding sequences of Clade J, additional sequences of *Miliolidium*, SymPortal database of named Defining Intragenomic Variants (DIVs, file “SymPortal_unique_DIVs” downloaded from symportal.org, last updated 2024-02-13), and sequences identified by SymPortal as Symbiodiniaceae during the run on the same data. Taxonomy was assigned separately to the dataset that was not curated by LULU. Genus assignments for Symbiodiniaceae in the curated dataset were checked for consistency with BLASTn hits. All Symbiodiniaceae ASVs had matches only from the Symbiodiniaceae family; the ASVs with BLASTn hits that did not correspond to the assigned genus were filtered out during the abundance filtration step, as their abundance did not exceed the thresholds. Labels indicating the closest identity match were assigned to ASVs with vsearch v2.14.1 [9] against SymPortal’s named database combined with *Miliolidium* and Clade J sequences with option “usearch_global”, and the hit with the highest identity value was selected as a label (multiple hits reported if several had the same shared identity). To reduce spurious ASVs and possible sample cross-contamination, ASVs with abundance that did not exceed 0.01% or 10 reads in any sample were filtered out, and abundances of remaining ASVs were set to 0 in the samples where they did not exceed these thresholds. Globally, the relative abundance threshold did not exclude any additional ASVs (or DIVs) compared to the minimal count threshold, and 111 out of 132 ASVs were kept in the filtered dataset. This filtered dataset was visualized with bar charts, heatmaps, and ordination measures; the unfiltered dataset was used to calculate alpha diversity indices.

As SymPortal is optimized for anthozoan samples (corals and sea anemones), we used it only to assign host profiles and confirm the robustness of conclusions. For the SymPortal run, files from different runs and aliquots were concatenated for each sample. Data was analysed with SymPortal [10], with the database populated by sequences from the SymPortal database that have a name associated with them and standard SymPortal reference sequences. Samples were analysed using only DIV assignment for all the samples (option “load”). For host samples, ITS2 profiles were assigned in a separate run without any additional abundance filtering (options “load” and “analyse”). Sequences derived from SymPortal (DIVs) and DADA2 (ASVs) were compared with vsearch to identify the degree of overlap.

Further analysis, including plot generation, was done with R 4.5.0 in RStudio 2024.12.1 [11]. All ASV and DIV sequences were aligned with MAFFT v7.505 [12] (option “auto”), the alignment was trimmed with trimAl v1.5 [13] (“automated1”), and a maximum likelihood tree was inferred with FastTree 2.1.11 [14] with the Generalized Time-Reversible model with CAT approximation, bootstrap supports were calculated by the default algorithm of FastTree as Shimodaira-Hasegawa (SH)-like local supports with 1000 resamples (Fig. S8). Trees were generated independently for each dataset (ASVs before LULU curation; ASVs after LULU curation; ASVs after LULU curation and filtration; DIVs; DIVs after filtration) to allow for more precision after removing rare highly divergent ASVs. We additionally tested two other approaches from [8], both creating a Neighbor-Joining tree based on a set of distances. The first approach is a hybrid tree based on between-genus distances approximated from the 28S rRNA Maximum Likelihood tree, and the intra-genus distances are calculated as pairwise distances between ITS2 sequences using identity. To get 28S distances, sequences of Clade J [15] were added to the previously published 28S alignment [16] with the MAFFT “add” option, then visually inspected and trimmed to the same length in Geneious Prime, and then the tree was built with the GTR+G+I model using IQ-TREE 2.4.0 [17]. The second approach is based on k-mer distances between ITS2 sequences (calculated with the kmer 1.1.2 package with k-mer size 7 and the EDGAR distance metric). Both methods gave comparable results but tended to distort between-genus relationships when highly divergent sequences were present, while FastTree preserved plausible genus relationships even on unfiltered data (Fig. S8). The k-mer tree emphasized variation between the sequences the most, while the tree generated by FastTree reduced this variation.

Diversity indices and Principal Coordinate Analysis (PCoA) plot based on weighted UniFrac distances were generated using the phyloseq 1.52.0 package [18]. Diversity indices were calculated on unfiltered data after LULU curation without rarefaction. Results with rarefaction to the minimum library depth (“rarefy_even_depth” in phyloseq) and with different filtration options (before and after LULU curation and filtering) were checked separately (Fig. S4). Main genera driving separation between sample types were visualized with distance-based redundancy analysis (db-RDA) using the “capscale” function in the vegan 2.7.1 [19] package (ordination constrained by sample type). The influence of sample type and site on the sequence composition was investigated with permutational multivariate analysis of variance (PERMANOVA) using the vegan package, and the pairwise differences between sample types were tested with the pairwiseAdonis 0.4.1 [20] package. Since the assumption of homogeneity of dispersions was not met in the full dataset (PERMDISP, *p* < 0.05), we excluded *Tridacna* samples, which were collected at a deeper part of the reef and displayed significantly greater within-group variability than most other sample types (TukeyHSD, *p* < 0.05 for multiple *Tridacna* comparisons, *p* > 0.05 for all non-*Tridacna* comparisons). For PCoA, PERMANOVA, and db-RDA, relative abundances of filtered, non-rarified data were used. ASV and DIV accumulation curves were built with the “specaccum” function in the vegan package.

The same analysis was performed on DIVs generated by SymPortal, with the following differences: no LULU curation; for PERMANOVA, both *Amphisorus* and *Tridacna* were removed as the presence of *Amphisorus* samples violated the assumption of homogeneity of dispersions (PERMDISP, *p* = 0.002).

R scripts for figure and table formatting and troubleshooting were assisted by ChatGPT (GPT-5, OpenAI) and subsequently manually reviewed, tested, and corrected by the authors.

All scripts used to analyse the data, the abundance table generated by DADA2 and SymPortal, and the database used to assign Symbiodiniaceae genera were uploaded to GitHub https://github.com/ECBSU/ECBSU_manuscripts_code/tree/main/Emelianenko_Comparing_Symbiodiniaceae_diversity-2026. Raw sequencing data were deposited in the NCBI Sequence Read Archive under the BioProject ID PRJNA1299766. Full SymPortal results were uploaded to FigShare 10.6084/m9.figshare.29881454.

### Host identification

Corals and Foraminifera were morphologically identified to the genus level during collection, and foraminiferal samples were additionally photographed in bright field with a Nikon DS-Ri2 camera mounted on an Olympus SZX16 stereomicroscope. To confirm host genus identity, we used molecular barcoding with the mitochondrial cytochrome c oxidase subunit I (COI) gene, mitochondrial 16S rRNA gene, 18S rRNA gene, and Internal Transcribed Spacer 1 (ITS1) of the rDNA region (Table S1). PCRs were performed in the volume of 25 μL, using 2–3 μL of template DNA and either OneTaq Hot Start 2X Master Mix with Standard Buffer (M048, New England Biolabs) or Q5 Hot Start High-Fidelity DNA Polymerase (M0493, New England Biolabs) according to the manufacturer’s protocol (Table S1). PCR products were visualized on a 1.5% agarose gel with Gel Red dye (Biotium 41002). PCR products were purified with the Monarch PCR & DNA Cleanup Kit (NEB) and sequenced using the SeqStudio Genetic Analyzer with BigDye Terminator v3.1 Cycle Sequencing Kit (Thermo Fisher Scientific) according to the manufacturer’s protocol and using the same primers as during amplification. Sequences for molecular barcoding were assembled and trimmed of primers and low-quality regions using Geneious Prime 2025.1.3. A check of the sequence identity was performed with BLASTn megablast against the nucleotide collection (nt) on the NCBI website (01-07-2025) [21], and sequences together with selected references were aligned with Muscle 5.1 [22] in Geneious Prime. Phylogenetic trees were inferred using IQ-TREE 2.4.0 [17] with the evolutionary model automatically selected by ModelFinder and 10,000 ultra-fast bootstrap replicates [23] and visualized with the ggtree 3.16.3 package [24]. The alignments and tree files have been uploaded to FigShare (10.6084/m9.figshare.29881454).

**Supplementary Table 1.**
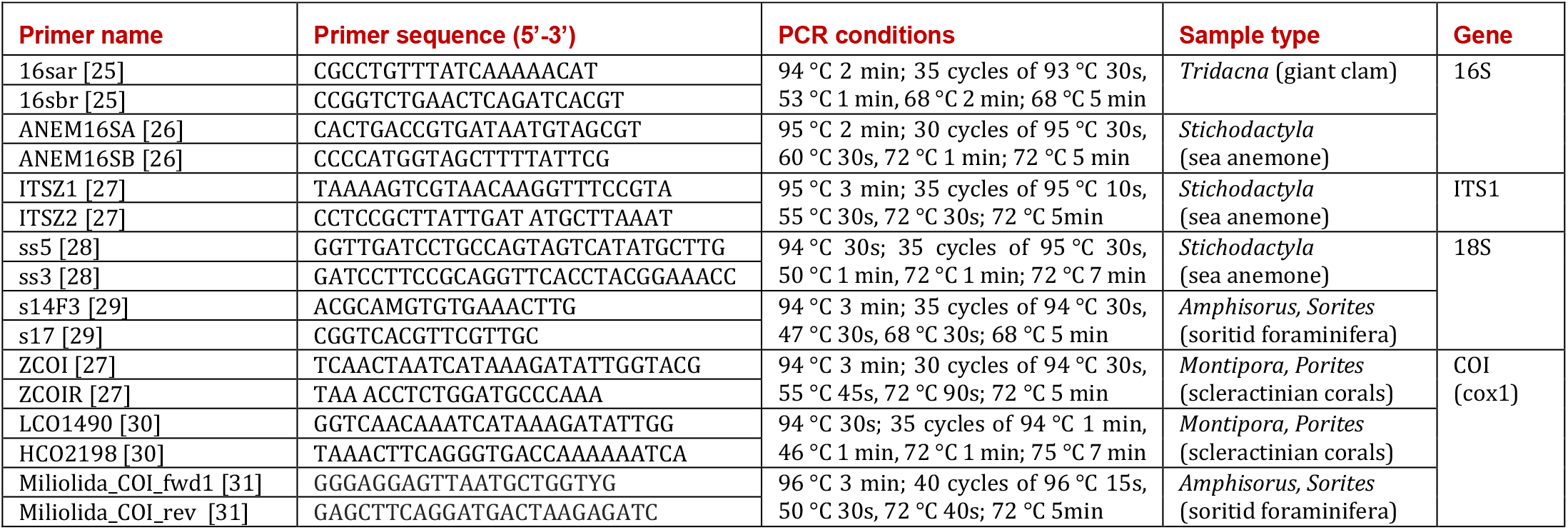
Primer sequences and PCR conditions used for host identification in the current study.

## Supplementary results and discussion

### ITS2 analysis

DADA2 and SymPortal produced an overlapping, but not identical, set of sequences: 97.8% of SymPortal DIVs had vsearch matches with more than 95% identity in DADA2 ASVs, but only 67.6% of ASVs had matches in DIVs with more than 95% identity (95% is the threshold used in LULU). Sequences from *Miliolidium* and Clade J were absent in SymPortal DIVs but reported as “non-Symbiodiniaceae sequences” instead. We primarily used DADA2 ASVs in downstream analysis, but our main conclusions were confirmed on SymPortal data (compare Fig. 1 and S3; Fig. 2 and S5; Tables S3–S5; Figs. S4, S6).

Metazoan hosts each contained 1-3 ASVs belonging to 1-2 genera: *Symbiodinium* and *Cladocopium* in *Tridacna*, and *Durusdinium* and *Cladocopium* in anthozoans (Fig. 1). *Symbiodinium* ITS2 variants A3 and A6 were detected exclusively in *Tridacna* samples. *Durusdinium* variants D1–D4 dominated most of the *Goniastrea* samples and were present in two *Porites* samples (two additional *Durusdinium* ASVs were present in one *Porites*). *Cladocopium* major ASVs were consistent with ITS2 profiles and differed between all genera of anthozoans, with C15 dominant in *Porites*, C21dp in *Montipora*, and C3gi-like ASV (98% identity) in *Stichodactyla*. C1 variant was detected in all giant clams and one *Goniastrea* sample, but was also present in all environmental and multiple foraminiferal samples. The C15 variant is predominantly found in *Porites* in the Indo-Pacific [32,33], and some C15 strains associated with shallow-water *Porites* were shown to exhibit high thermal tolerance [34]. *Montipora* are known to host variants related to C21 in the Indo-Pacific [33,35]. *Stichodactyla* sea anemones around Okinawa were shown to associate with *Cladocopium*, although no ITS2 type was specified [36]. ITS2 type C1 has also been previously reported from Indo-Pacific *Tridacna maxima* [35], *Goniastrea* [35], and *Amphisorus* [37].

Foraminiferal symbionts were genus-specific apart from *Cladocopium. Amphisorus* samples were strongly dominated by *Freudenthalidium* (F3w and 2 ASVs, both ∼96-97% identical to F3o) or *Cladocopium. Sorites* samples contained *Fugacium* (F5.1, F51a), *Miliolidium* (91% identity to D1.2), and *Cladocopium*. While *Cladocopium* ITS2 profiles were different in Foraminifera (Fig. 1D), abundant profiles consistently had C1 as the main ASV (Fig. 1C). Low proportions of *Durusdinium* and *Symbiodinium* were also detected in some foraminiferal samples.

Foraminiferal and environmental samples contained *Fugacium, Freudenthalidium*, and *Miliolidium*, which is consistent with these genera being considered primarily foraminiferal symbionts or free-living [16,38,39]. *Miliolidium* was also present in most macroalgal and half of the sediment samples, indicating potentially more soritid hosts or free-living *Miliolidium. Miliolidium* ITS2 sequences exceeded 310 bp and were not identified by SymPortal (as well as Clade J), which may explain why it is rarely reported from sediments [40]. Overlap between benthic environmental and foraminiferal samples can be explained by the presence of small Foraminifera in the environmental samples, expelled Symbiodiniaceae, free-living stages of symbionts, the same ITS2 variants shared among multiple Symbiodiniaceae species, and Foraminifera ingesting Symbiodiniaceae as food. The absence of *Halluxium* in our *Amphisorus* samples is consistent with recent work showing that shallow water *Amphisorus* in Okinawa associate with *Freudenthalidium* and *Cladocopium*, while *Halluxium* is only found in deeper dwelling *Amphisorus* [41]. Overall, the presence of *Freudenthalidium, Fugacium, Miliolidium, Halluxium*, Clades I–J, and Fr4 primarily or exclusively in macroalgal and sediment samples is consistent with their benthic niche.

Environmental Symbiodiniaceae communities vary at both global and local spatial, as well as temporal, scales [8,42,43], and sample processing differs among studies. Although previous work shows that even a small spatial change can result in different Symbiodiniaceae composition in environmental samples, we could not detect a significant effect of the site (PERMANOVA limited to environmental samples, p > 0.05); a weak effect was detected only for water samples when analysed with SymPortal data (PERMANOVA, p = 0.032). We also could not detect any difference between macroalgal samples consisting of *Padina* sp. (Fig. 1B, columns 1, 4, 5) and miscellaneous macroalgae (the other macroalgal samples) (PERMANOVA, p = 0.669). While we sampled the reef flat only once, it would be interesting to see the temporal dynamics of the Symbiodiniaceae community in the same place as it is known to shift depending on the season [8].

We chose ITS2 because it is one of the most widely used markers for Symbiodiniaceae, thus allowing for greater comparability with previous studies, and its high diversity allows for fine-scale intra-genus resolution. However, diversity metrics obtained with this marker should be interpreted with caution. ITS2 data does not directly enable species-level resolution: the same ITS2 variant may be present within different Symbiodiniaceae species or exhibit intra-specific variability [44]. We applied SymPortal profiles and LULU curation to mitigate the intra-species variability. However, it is impossible to use SymPortal profiles on the environmental data, as the central assumption of SymPortal of one Symbiodiniaceae species per genus per host is violated in the environment and potentially in some non-coral hosts. At the same time, LULU curation would fail to merge some variants that are coming from the same cell if they are beyond similarity cut-off (D1 and D4) or, on the contrary, would merge some variants that might belong to different species but are co-occurring for ecological reasons (for example, several species of *Symbiodinium* specific to only water samples because this is their environmental niche).

Relative abundances of ITS2 sequences should also be interpreted with caution, as they might not accurately represent relative abundances of corresponding genera, because the ITS2 region is known to have considerable, sometimes 10-fold, copy number variation between Symbiodiniaceae genera [44,45]. For example, in foraminiferal samples, non-Symbiodiniaceae ASVs and low-abundance genera were primarily present in *Sorites* (dominated by *Fugacium*) and *Freudenthalidium*-dominated *Amphisorus*, while completely absent in *Cladocopium*-dominated *Amphisorus*, even in unfiltered data (Fig. 1B–C, Fig. S1). This could be explained by higher ITS2 copy number in *Cladocopium* (known to exceed 2000 copies per cell in some species [45]) masking low-abundance ASVs, which could represent true background symbionts, biological contamination (recent ingestion of free-living cells), or a mixture of both. Correspondingly, *Amphisorus* samples in Fig. 2C demonstrate a bimodal distribution of diversity metrics with *Cladocopium*-dominated samples at the bottom (no other ASVs) and *Freudenthalidium*-dominated samples at the top.

### Host identification

Marker gene sequences obtained for host samples are presented in Table S6. Two samples of *Sorites* (1-9 and 1-12) were not barcoded due to the lack of available DNA after the library preparation, and for Sorites2-21 and Sorites2-29, only the partial COI gene was sequenced. *Goniastrea* samples were not barcoded because no visible PCR products were obtained with either the universal COI primers LCO1490–HCO2198 or the coral-specific primers ZCOI– ZCOIR. Attempts to use the coral-specific ribosomal internal transcribed spacer region (ITS) with primers ITSZ1 and ITSZ2 for molecular barcoding resulted in amplification of Symbiodiniaceae instead of the host. When amplified with foraminifera-specific primers, sample Goniastrea1-3 resulted in the amplification of COI with 90% identity to *Borelis* (Alveolinidae, Foraminifera) (Fig. S9). However, since this genus of foraminifera is known to harbor only diatom symbionts, its presence cannot explain the different Symbiodiniaceae genus in this sample and probably represents contamination. Additionally, COI amplification has failed for *Porites* samples Porites2-3 and Porites2-7. However, as anthozoan hosts were not the focus of our study, and because there were no differences in symbiont composition among most *Goniastrea* samples (except Goniastrea1-3) and *Porites* samples, we did not continue attempts to barcode the hosts.

Soritid foraminifera exhibit high morphological variability and distinct genotypes [46]; therefore, we refrain from assigning our specimens to a specific species and provide marker gene sequences instead (Table S6). All the barcoded *Sorites* samples showed the same COI and 18S sequences, apart from Sorites1-13 (dominated by *Miliolidium*), which had 99.4% similarity in partial COI (324 bp) and 93.6% similarity in partial 18S rRNA gene sequences (324 bp) to the sequences derived from other samples. COI sequences of *Amphisorus* were identical but for one nucleotide (“C” in 1-4, 1-5, 1-10, 2-2, 2-6, 2-31; “T” in 1-3, 2-3, 2-5; not sequenced in 2-30, 1-7, 2-1). Within *Amphisorus*, two partial 18S rRNA gene variants were detected, consistent with COI difference and mostly associated with different symbionts, but not a specific site (Fig. S9). The first 18S rRNA gene variant, 100% identical to *Amphisorus kudakajimaensis* (LC813322), was detected in samples 1-2, 1-3, 1-7, 2-1, 2-3, 2-5, all of which, except 2-5, were associated with *Freudenthalidium*. The second 18S rRNA gene variant, sharing ∼92% identity with the first one, was detected in samples 1-4, 1-5, 1-10, 2-2, 2-4, 2-6, 2-30, 2-31, all of which, except 2-6, were associated with *Cladocopium*. Unlike most *Amphisorus* samples presented in [41], which showed comparable proportions of both symbionts in most samples, all of our *Amphisorus* samples were strongly dominated by one symbiont genus. It has been shown that in some locations, *Freudenthalidium* and *Cladocopium* C1 associate with genetically distinct *Amphisorus* [37]. Although we could also detect two *Amphisorus* genotypes primarily hosting distinct symbionts (Fig. S9), our limited sample size and sampling location do not allow us to determine whether symbionts are indeed host-specific within *Amphisorus*. Since all the samples were collected on one reef flat, even though the sites were separated by ∼200 m, we cannot exclude relatedness among the sampled Foraminifera, which could explain the symbiont distribution if the symbionts were transmitted vertically during asexual reproduction.

**Supplementary Table 2.**
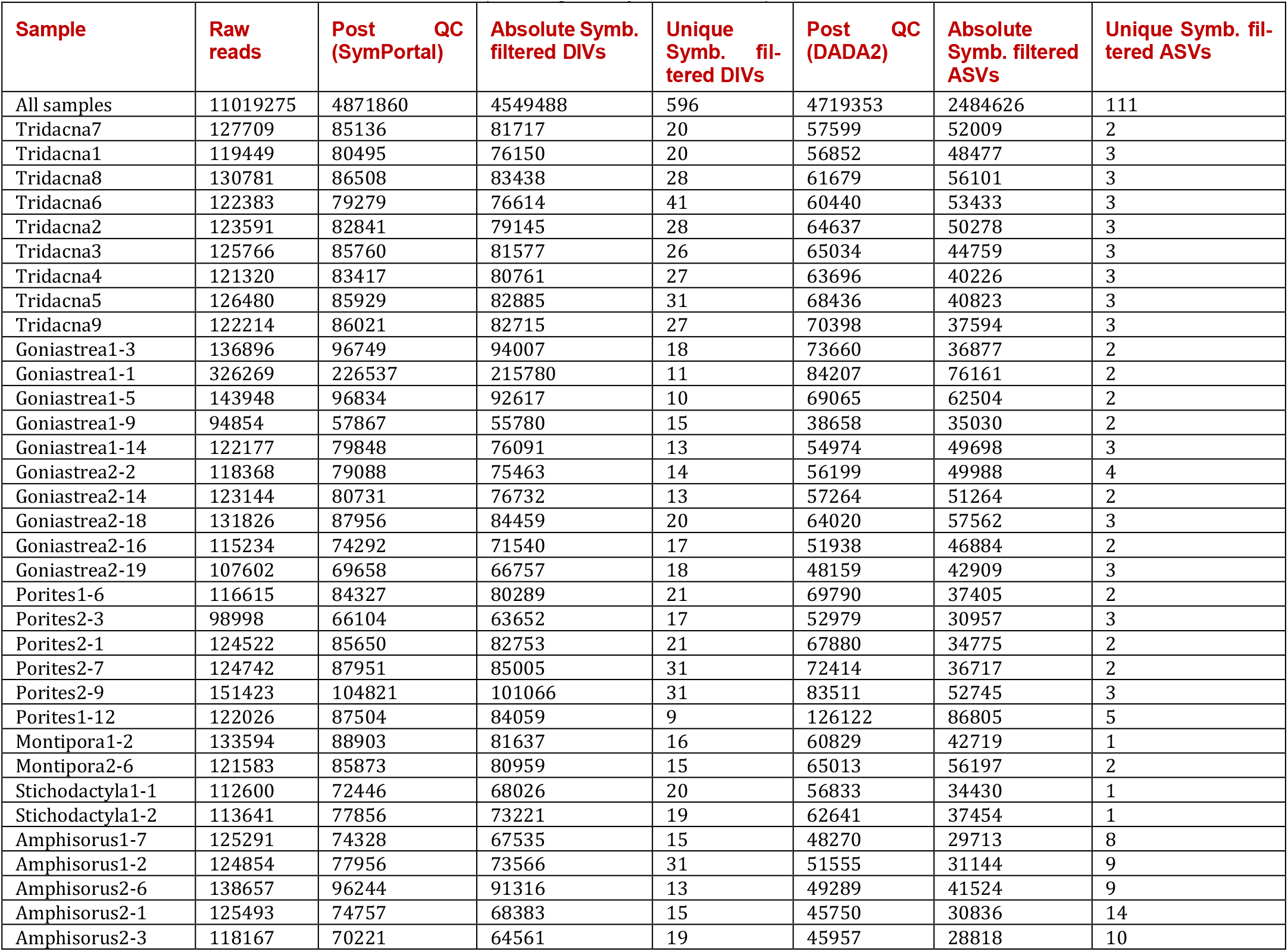
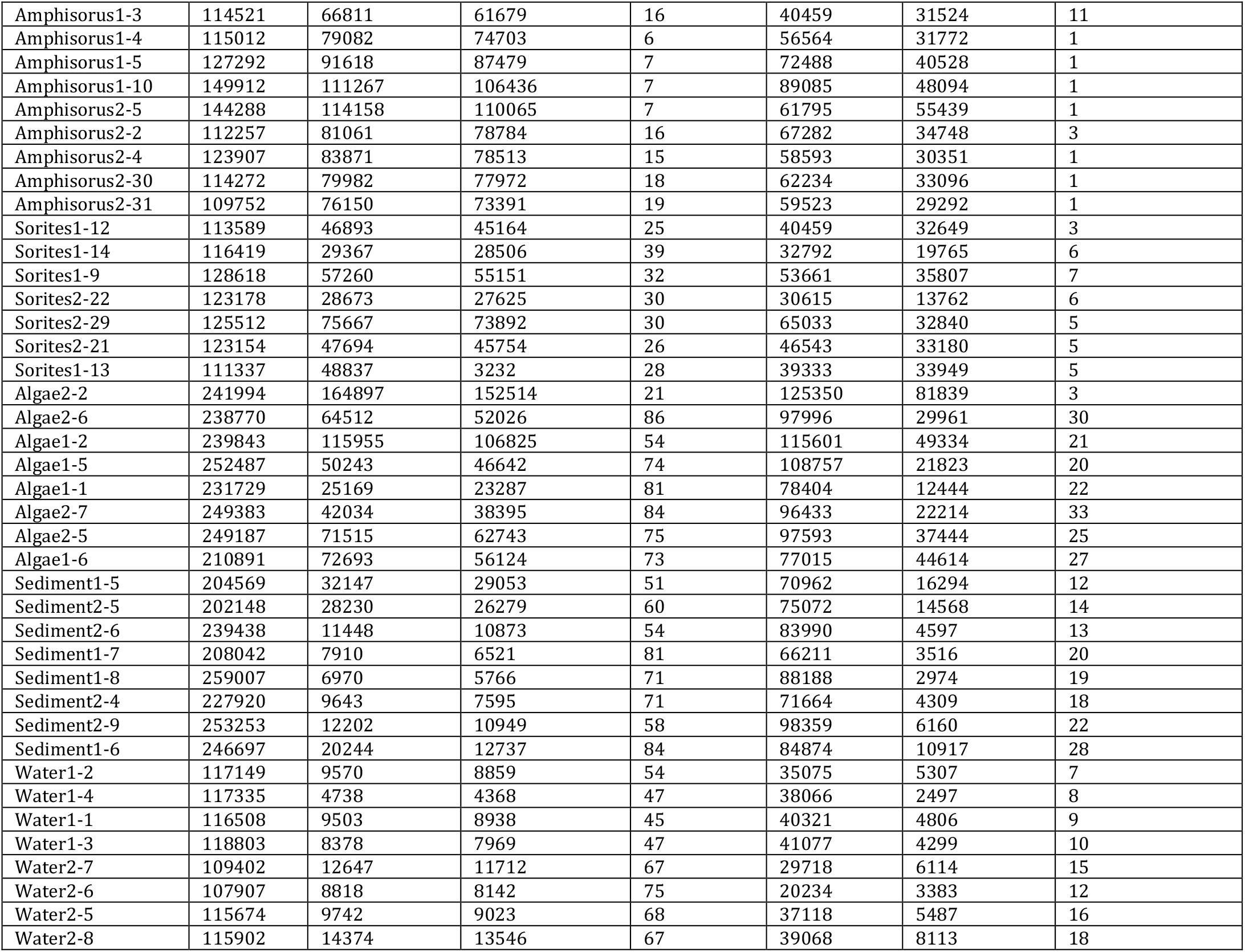
Summary of the number of reads in each sample before and after filtering by both SymPortal and DADA2. All samples had a total DIV count of 4549648 (721 unique DIVs) and Symbiodiniaceae ASV count of 2484626 (132 unique) before filtration, which excluded rare, low-abundance ASVs. Total ASV count (including non-Symbiodiniaceae) was 3060114.

| Sample | Raw reads | Post QC (SymPortal) | Absolute Symb. filtered DIVs | Unique Symb. filtered DIVs | Post QC (DADA2) | Absolute Symb. filtered ASVs | Unique Symb. filtered ASVs |
| --- | --- | --- | --- | --- | --- | --- | --- |
| All samples | 11019275 | 4871860 | 4549488 | 596 | 4719353 | 2484626 | 111 |
| *Tridacna*7 | 127709 | 85136 | 81717 | 20 | 57599 | 52009 | 2 |
| *Tridacna*1 | 119449 | 80495 | 76150 | 20 | 56852 | 48477 | 3 |
| *Tridacna*8 | 130781 | 86508 | 83438 | 28 | 61679 | 56101 | 3 |
| *Tridacna*6 | 122383 | 79279 | 76614 | 41 | 60440 | 53433 | 3 |
| *Tridacna*2 | 123591 | 82841 | 79145 | 28 | 64637 | 50278 | 3 |
| *Tridacna*3 | 125766 | 85760 | 81577 | 26 | 65034 | 44759 | 3 |
| *Tridacna*4 | 121320 | 83417 | 80761 | 27 | 63696 | 40226 | 3 |
| *Tridacna*5 | 126480 | 85929 | 82885 | 31 | 68436 | 40823 | 3 |
| *Tridacna*9 | 122214 | 86021 | 82715 | 27 | 70398 | 37594 | 3 |
| *Goniastrea*1-3 | 136896 | 96749 | 94007 | 18 | 73660 | 36877 | 2 |
| *Goniastrea*1-1 | 326269 | 226537 | 215780 | 11 | 84207 | 76161 | 2 |
| *Goniastrea*1-5 | 143948 | 96834 | 92617 | 10 | 69065 | 62504 | 2 |
| *Goniastrea*1-9 | 94854 | 57867 | 55780 | 15 | 38658 | 35030 | 2 |
| *Goniastrea*1-14 | 122177 | 79848 | 76091 | 13 | 54974 | 49698 | 3 |
| *Goniastrea*2-2 | 118368 | 79088 | 75463 | 14 | 56199 | 49988 | 4 |
| *Goniastrea*2-14 | 123144 | 80731 | 76732 | 13 | 57264 | 51264 | 2 |
| *Goniastrea*2-18 | 131826 | 87956 | 84459 | 20 | 64020 | 57562 | 3 |
| *Goniastrea*2-16 | 115234 | 74292 | 71540 | 17 | 51938 | 46884 | 2 |
| *Goniastrea*2-19 | 107602 | 69658 | 66757 | 18 | 48159 | 42909 | 3 |
| *Porites*1-6 | 116615 | 84327 | 80289 | 21 | 69790 | 37405 | 2 |
| *Porites*2-3 | 98998 | 66104 | 63652 | 17 | 52979 | 30957 | 3 |
| *Porites*2-1 | 124522 | 85650 | 82753 | 21 | 67880 | 34775 | 2 |
| *Porites*2-7 | 124742 | 87951 | 85005 | 31 | 72414 | 36717 | 2 |
| *Porites*2-9 | 151423 | 104821 | 101066 | 31 | 83511 | 52745 | 3 |
| *Porites*1-12 | 122026 | 87504 | 84059 | 9 | 126122 | 86805 | 5 |
| *Montipora*1-2 | 133594 | 88903 | 81637 | 16 | 60829 | 42719 | 1 |
| *Montipora*2-6 | 121583 | 85873 | 80959 | 15 | 65013 | 56197 | 2 |
| *Stichodactyla*1-1 | 112600 | 72446 | 68026 | 20 | 56833 | 34430 | 1 |
| *Stichodactyla*1-2 | 113641 | 77856 | 73221 | 19 | 62641 | 37454 | 1 |
| *Amphisorus*1-7 | 125291 | 74328 | 67535 | 15 | 48270 | 29713 | 8 |
| *Amphisorus*1-2 | 124854 | 77956 | 73566 | 31 | 51555 | 31144 | 9 |
| *Amphisorus*2-6 | 138657 | 96244 | 91316 | 13 | 49289 | 41524 | 9 |
| *Amphisorus*2-1 | 125493 | 74757 | 68383 | 15 | 45750 | 30836 | 14 |
| *Amphisorus*2-3 | 118167 | 70221 | 64561 | 19 | 45957 | 28818 | 10 |
| Amphisorus1-3 | 114521 | 66811 | 61679 | 16 | 40459 | 31524 | 11 |
| Amphisorus1-4 | 115012 | 79082 | 74703 | 6 | 56564 | 31772 | 1 |
| Amphisorus1-5 | 127292 | 91618 | 87479 | 7 | 72488 | 40528 | 1 |
| Amphisorus1-10 | 149912 | 111267 | 106436 | 7 | 89085 | 48094 | 1 |
| Amphisorus2-5 | 144288 | 114158 | 110065 | 7 | 61795 | 55439 | 1 |
| Amphisorus2-2 | 112257 | 81061 | 78784 | 16 | 67282 | 34748 | 3 |
| Amphisorus2-4 | 123907 | 83871 | 78513 | 15 | 58593 | 30351 | 1 |
| Amphisorus2-30 | 114272 | 79982 | 77972 | 18 | 62234 | 33096 | 1 |
| Amphisorus2-31 | 109752 | 76150 | 73391 | 19 | 59523 | 29292 | 1 |
| Sorites1-12 | 113589 | 46893 | 45164 | 25 | 40459 | 32649 | 3 |
| Sorites1-14 | 116419 | 29367 | 28506 | 39 | 32792 | 19765 | 6 |
| Sorites1-9 | 128618 | 57260 | 55151 | 32 | 53661 | 35807 | 7 |
| Sorites2-22 | 123178 | 28673 | 27625 | 30 | 30615 | 13762 | 6 |
| Sorites2-29 | 125512 | 75667 | 73892 | 30 | 65033 | 32840 | 5 |
| Sorites2-21 | 123154 | 47694 | 45754 | 26 | 46543 | 33180 | 5 |
| Sorites1-13 | 111337 | 48837 | 3232 | 28 | 39333 | 33949 | 5 |
| Algae2-2 | 241994 | 164897 | 152514 | 21 | 125350 | 81839 | 3 |
| Algae2-6 | 238770 | 64512 | 52026 | 86 | 97996 | 29961 | 30 |
| Algae1-2 | 239843 | 115955 | 106825 | 54 | 115601 | 49334 | 21 |
| Algae1-5 | 252487 | 50243 | 46642 | 74 | 108757 | 21823 | 20 |
| Algae1-1 | 231729 | 25169 | 23287 | 81 | 78404 | 12444 | 22 |
| Algae2-7 | 249383 | 42034 | 38395 | 84 | 96433 | 22214 | 33 |
| Algae2-5 | 249187 | 71515 | 62743 | 75 | 97593 | 37444 | 25 |
| Algae1-6 | 210891 | 72693 | 56124 | 73 | 77015 | 44614 | 27 |
| Sediment1-5 | 204569 | 32147 | 29053 | 51 | 70962 | 16294 | 12 |
| Sediment2-5 | 202148 | 28230 | 26279 | 60 | 75072 | 14568 | 14 |
| Sediment2-6 | 239438 | 11448 | 10873 | 54 | 83990 | 4597 | 13 |
| Sediment1-7 | 208042 | 7910 | 6521 | 81 | 66211 | 3516 | 20 |
| Sediment1-8 | 259007 | 6970 | 5766 | 71 | 88188 | 2974 | 19 |
| Sediment2-4 | 227920 | 9643 | 7595 | 71 | 71664 | 4309 | 18 |
| Sediment2-9 | 253253 | 12202 | 10949 | 58 | 98359 | 6160 | 22 |
| Sediment1-6 | 246697 | 20244 | 12737 | 84 | 84874 | 10917 | 28 |
| Water1-2 | 117149 | 9570 | 8859 | 54 | 35075 | 5307 | 7 |
| Water1-4 | 117335 | 4738 | 4368 | 47 | 38066 | 2497 | 8 |
| Water1-1 | 116508 | 9503 | 8938 | 45 | 40321 | 4806 | 9 |
| Water1-3 | 118803 | 8378 | 7969 | 47 | 41077 | 4299 | 10 |
| Water2-7 | 109402 | 12647 | 11712 | 67 | 29718 | 6114 | 15 |
| Water2-6 | 107907 | 8818 | 8142 | 75 | 20234 | 3383 | 12 |
| Water2-5 | 115674 | 9742 | 9023 | 68 | 37118 | 5487 | 16 |
| Water2-8 | 115902 | 14374 | 13546 | 67 | 39068 | 8113 | 18 |

**Supplementary Figure 1.**
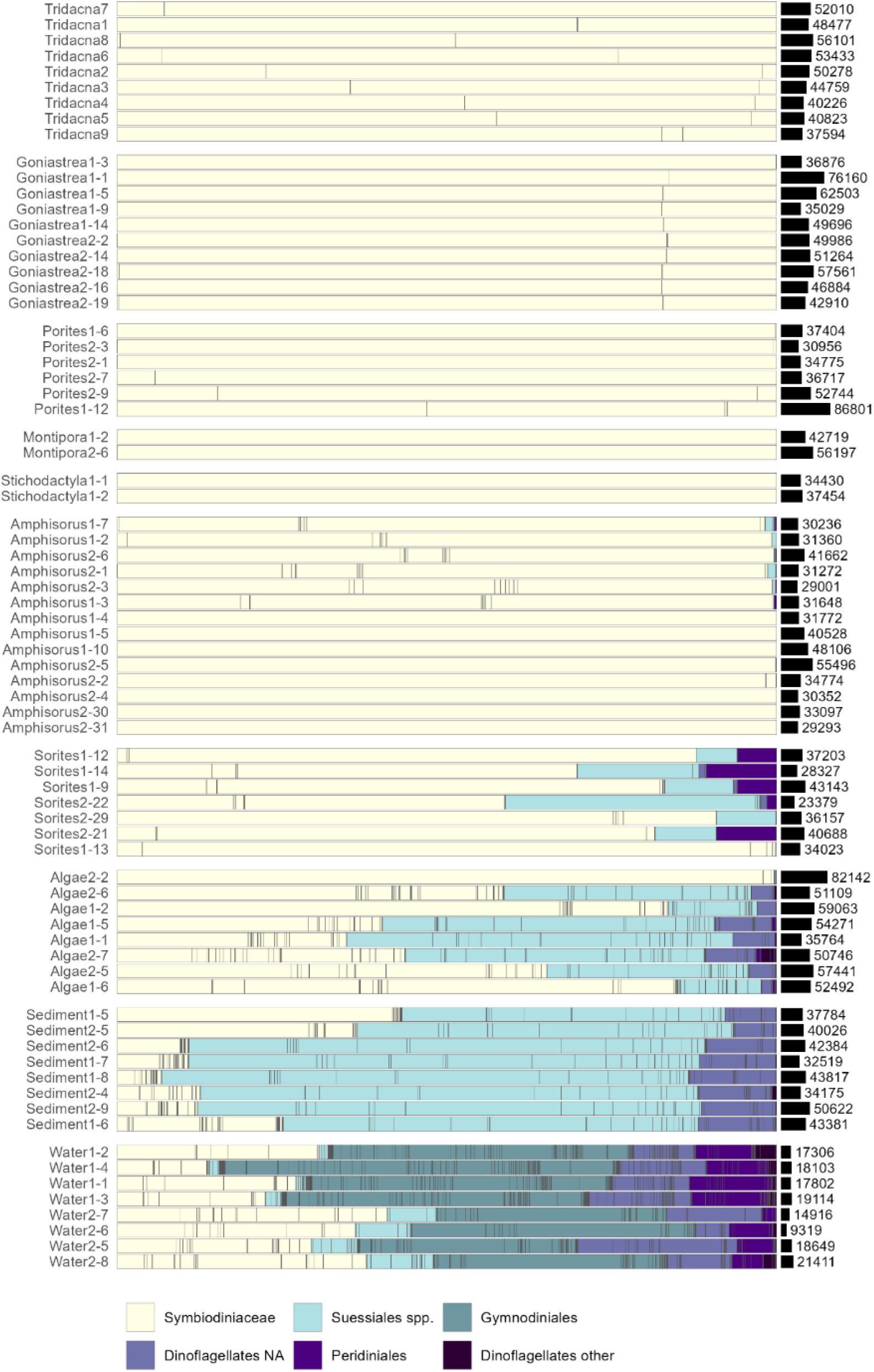
Sample composition according to DADA2 taxonomic annotation showing all dinoflagellate ASVs, annotated based on the DADA2 pipeline (the non-dinoflagellate ASVs were filtered out). The grey lines separate individual ASVs. “Suessiales spp.” refers to non-Symbiodiniaceae Suessiales regardless of their genus annotation. “Dinoflagellates other” include orders Prorocentrales, Thoracosphaerales, and Dinophysiales. The small black bars and numbers on the right represent the total number of reads per sample. The sediment samples demonstrate relatively high dinoflagellate ASV counts but a small number of Symbiodiniaceae DIVs, which can be explained by non-target amplification of other dinoflagellates, mostly other Suessiales. Water samples also have many non-target ASVs (>50%) and fewer reads per sample, as they were not merged from different aliquots. The non-Symbiodiniaceae Suessiales in *Sorites* samples is represented by ASV16, having low sequence identity to *Pelagodinium* (∼75% over ∼310 bp). This ASV is also present in samples of *Amphisorus*, macroalgae, and sediment. The Peridiniales in *Sorites* are represented mainly by ASV36, which matches *Vulcanodinium rugosum* with 99.68% identity and 100% query coverage. Sample Algae2-2 has no non-target amplification and was removed from the downstream analysis.

**Supplementary Figure 2.**
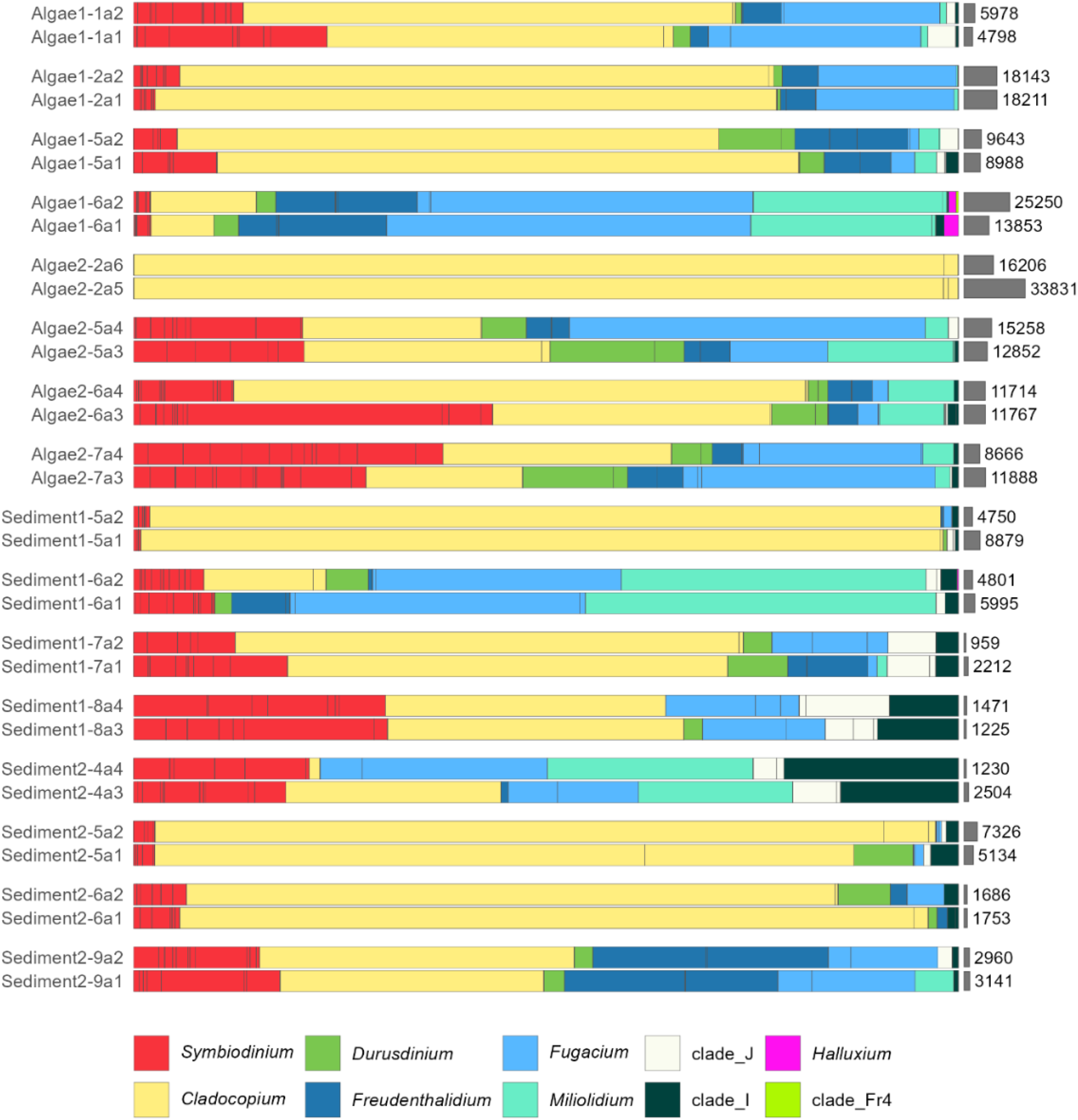
Relative abundance of Symbiodiniaceae DIVs in different aliquots from the macroalgae and sediment bulk samples. Numbers and grey bars on the right represent total counts (abundance) of Symbiodiniaceae DIVs. The grey lines separate individual ASVs. Sample Algae2-2 has no non-*Cladocopium* ASVs or non-target dinoflagellate amplification and was removed from further analysis as an outlier. It is possible that this sample contained a piece of host (one or several *Amphisorus*), whose symbionts masked other Symbiodiniaceae genera due to high ITS2 copy number [44,45]. Clade Fr4 is only present in one of the samples (Algae1-6a2).

**Supplementary Figure 3.**
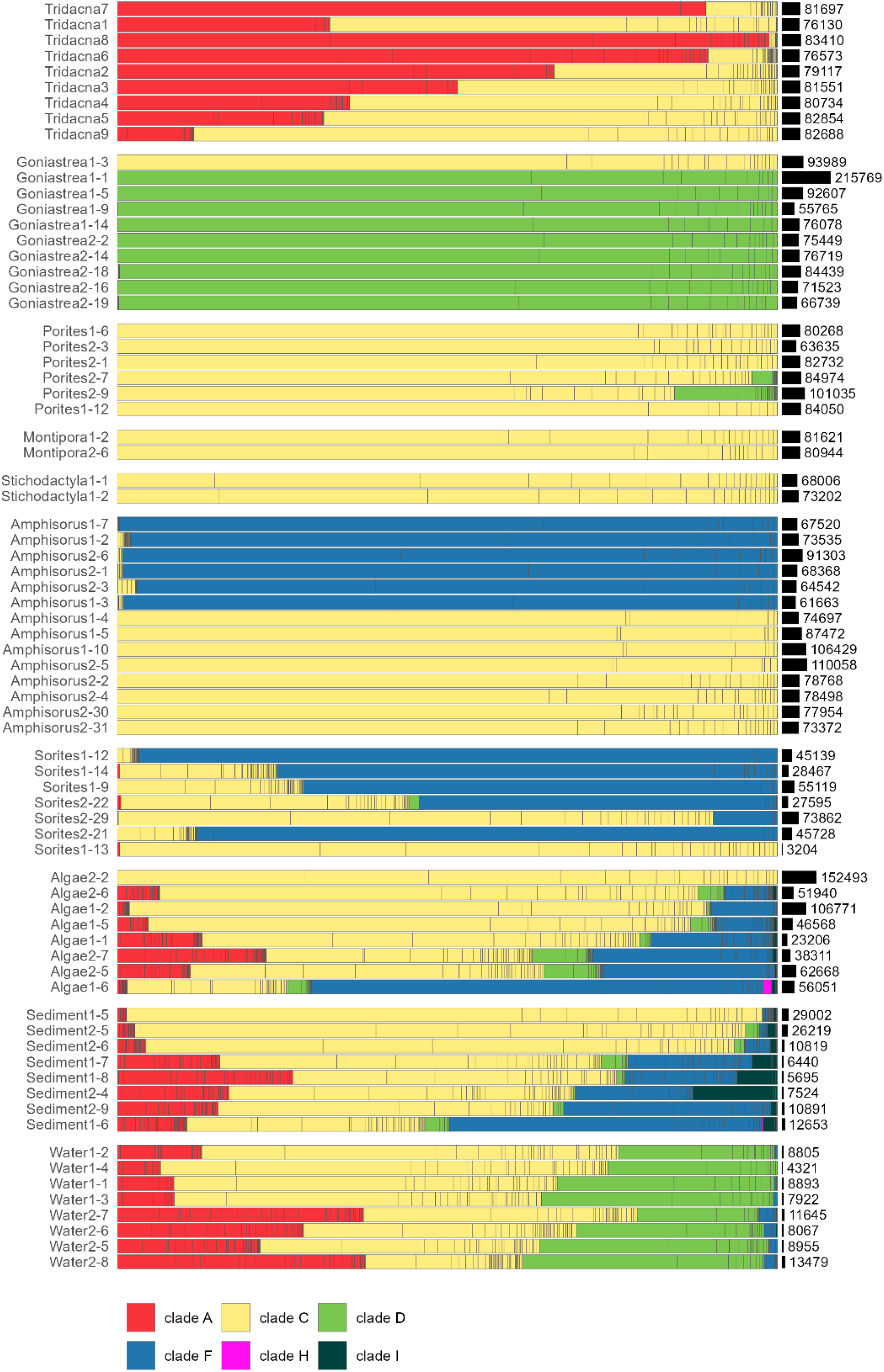
Sample composition according to SymPortal. The small black bars and numbers on the right represent the total number of reads per sample. A few reads (<22) were assigned to *Breviolum* but filtered out during the relative abundance / minimal count filtration step. Main differences include the absence of Clade J, *Miliolidium* (in Sorites1-13), and no genus division of Clade F (compare to Fig.1A). Sequences of Clade J and *Miliolidium* were identified but not quantified by SymPortal as they were classified as “non-Symbiodiniaceae sequences”.

**Supplementary Figure 4.**
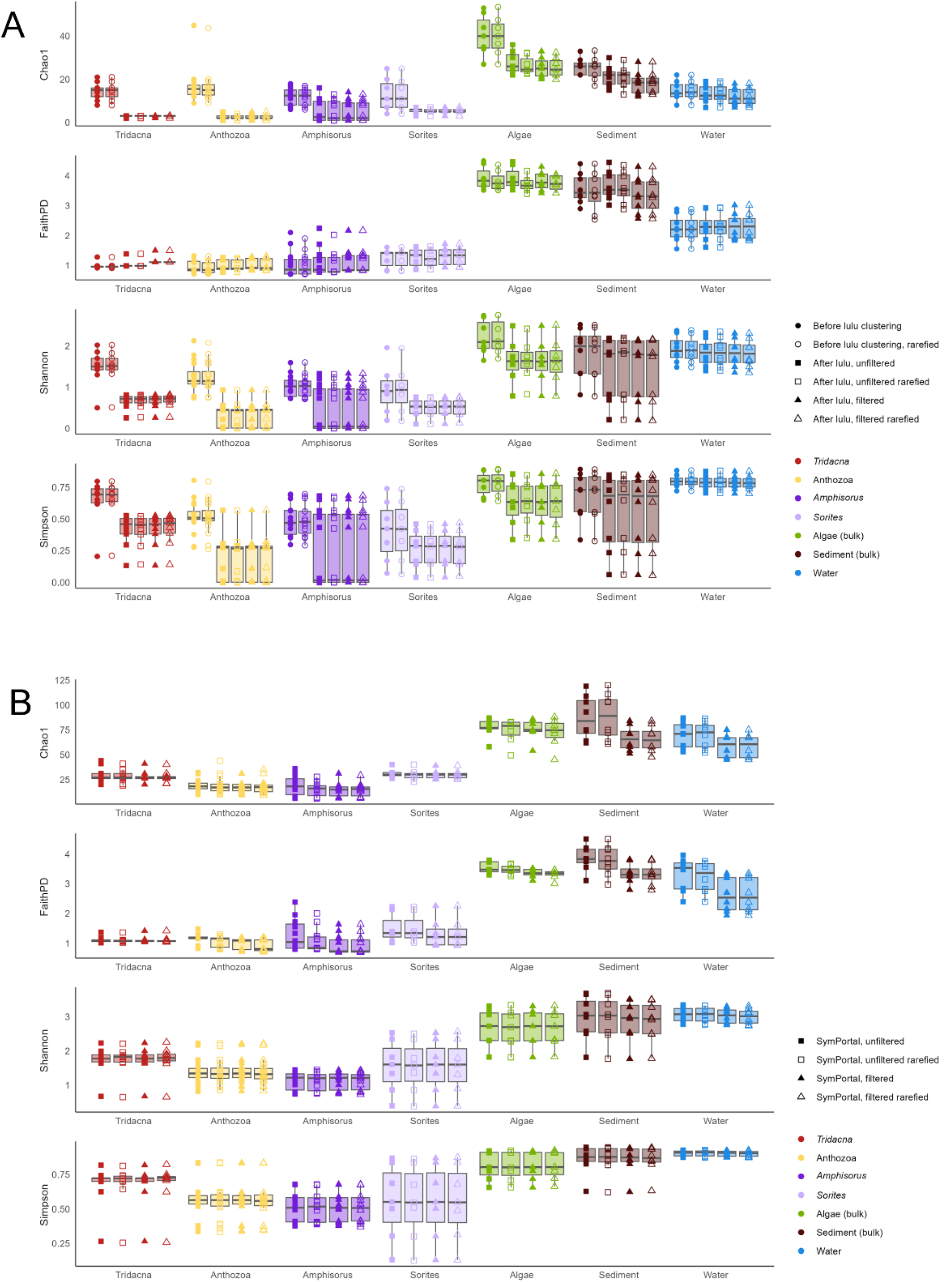
Comparison between the alpha diversity measures across different clustering and filtering parameters and rarefaction. A – Based on data from the DADA2+LULU pipeline, B – Based on data from the SymPortal. Samples were rarified to the lowest sequencing depth: 2489 reads for the filtered dataset and 2511 for the unfiltered dataset in the case of the DADA2+LULU pipeline, 3204 reads for SymPortal. LULU processing caused significant reductions in the diversity measures (median percent change across samples: Chao1 −70%, Shannon −52%, Simpson −41%; Faith’s PD 3%), whereas subsequent filtering and rarefaction had negligible effects on ASV data. Chao1 in filtered samples equals richness.

**Supplementary Figure 5.**
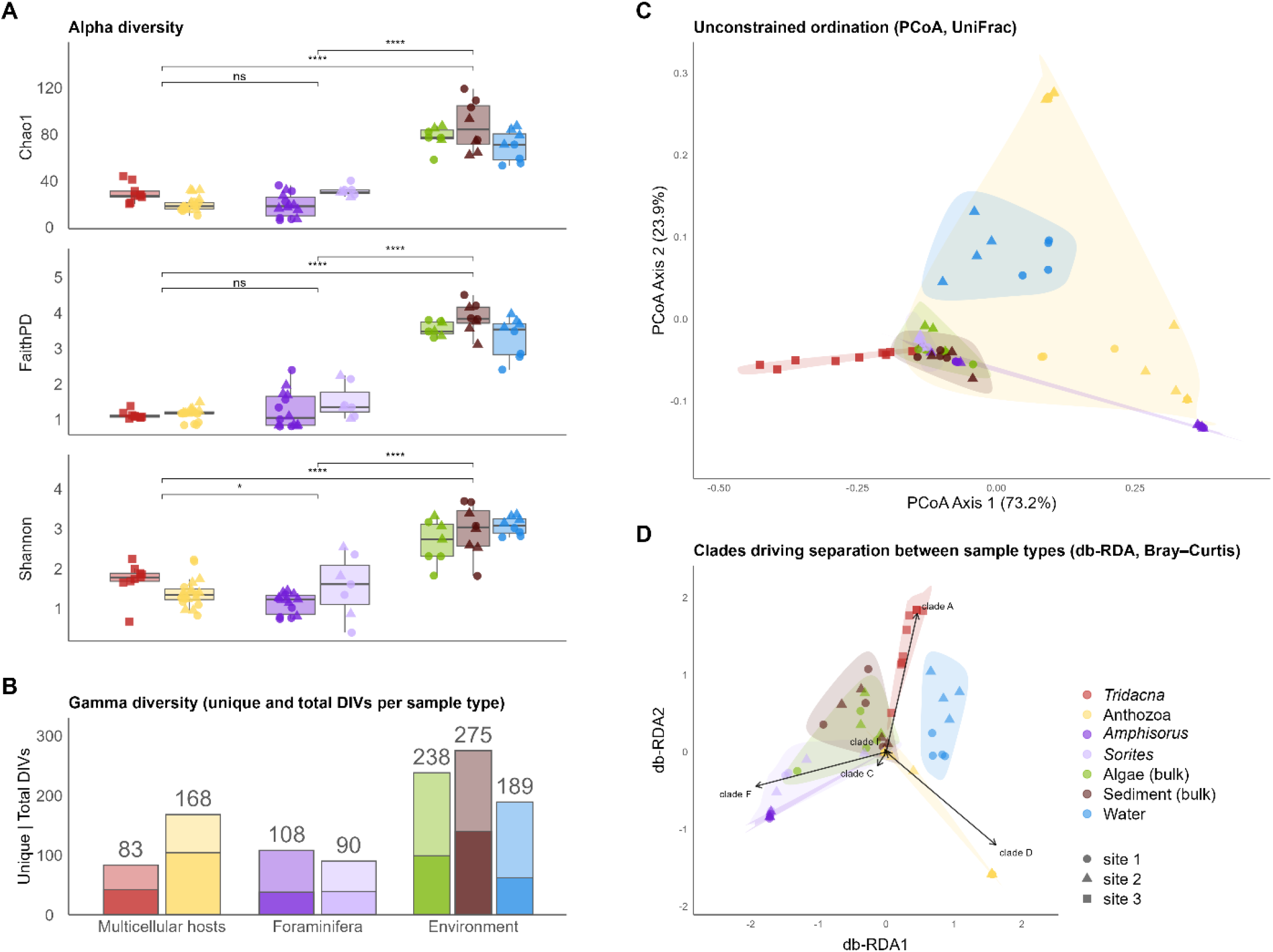
Diversity measures and comparison between the habitats based on SymPortal data. The plots are analogous to Fig. 2.

**Supplementary Figure 6.**
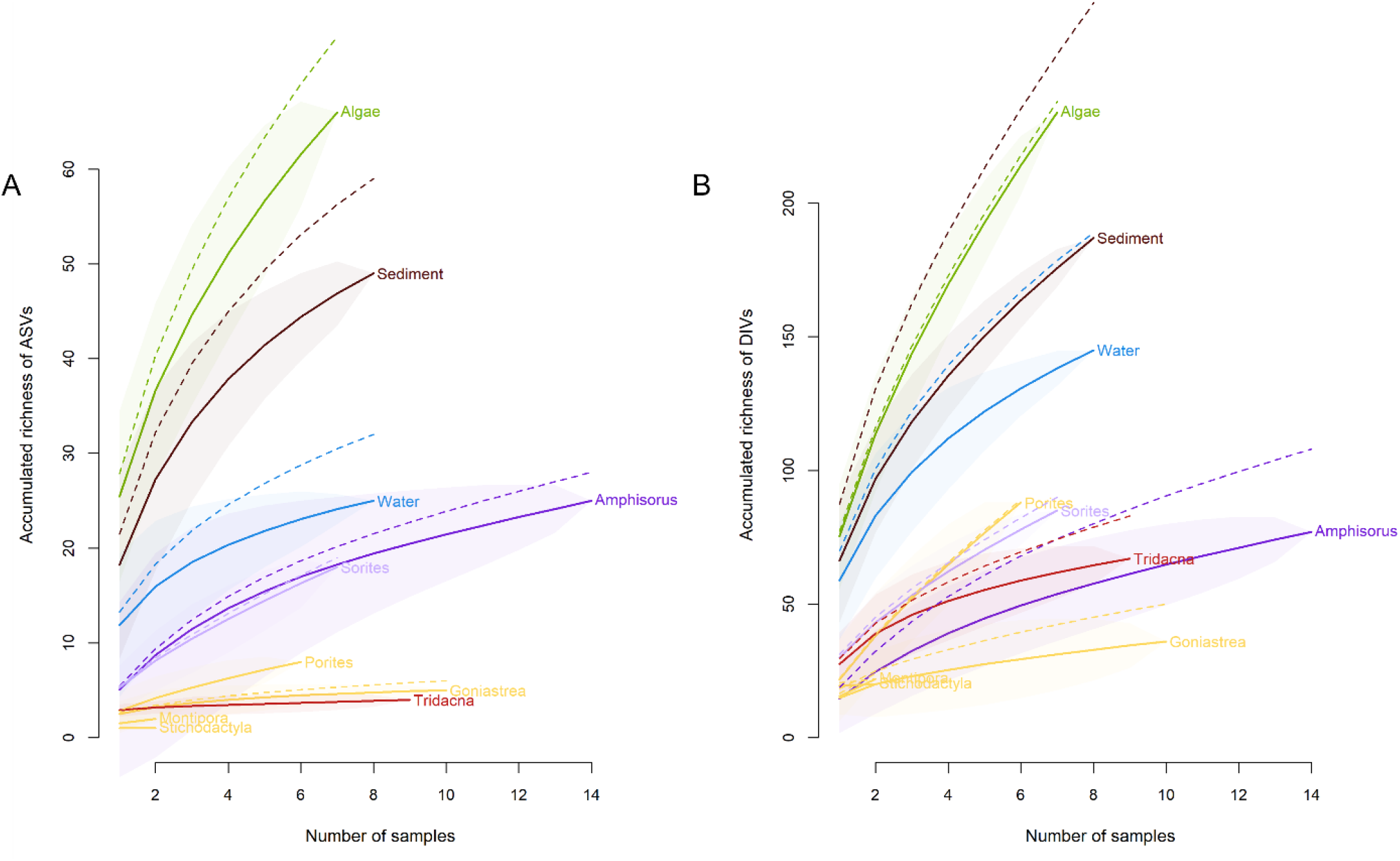
Richness accumulation curves. A – Based on ASVs (DADA2+LULU pipeline). B – Based on DIVs (SymPortal). Unfiltered data are represented by dotted lines and filtered data by solid lines with confidence intervals.

**Supplementary Figure 7.**
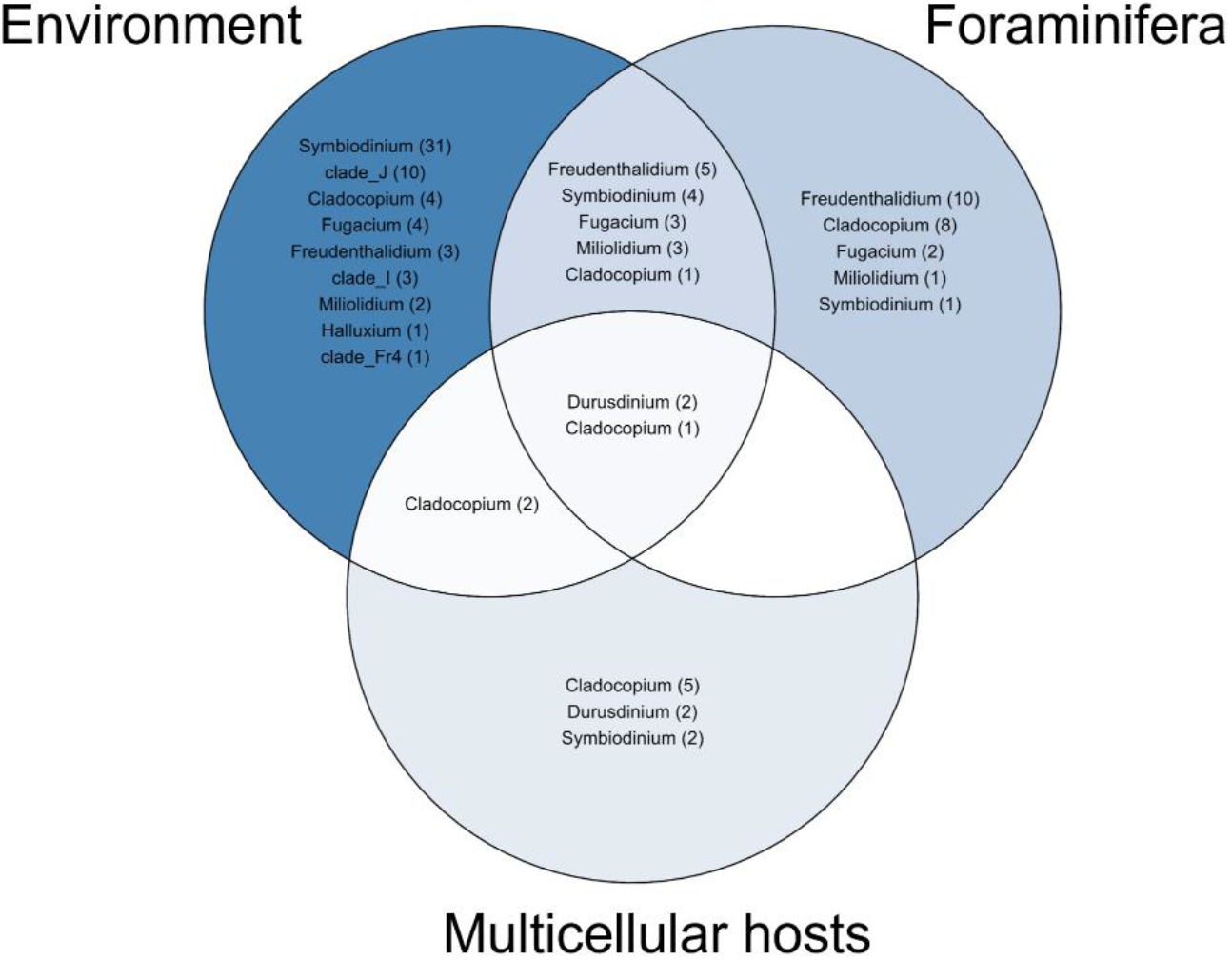
A Venn diagram showing ASV overlaps among three sample categories. Based on the DADA2+LULU pipeline, filtered dataset. Darker shades represent a higher number of sequences, and numbers in brackets represent the number of ASVs.

**Supplementary Figure 8.**
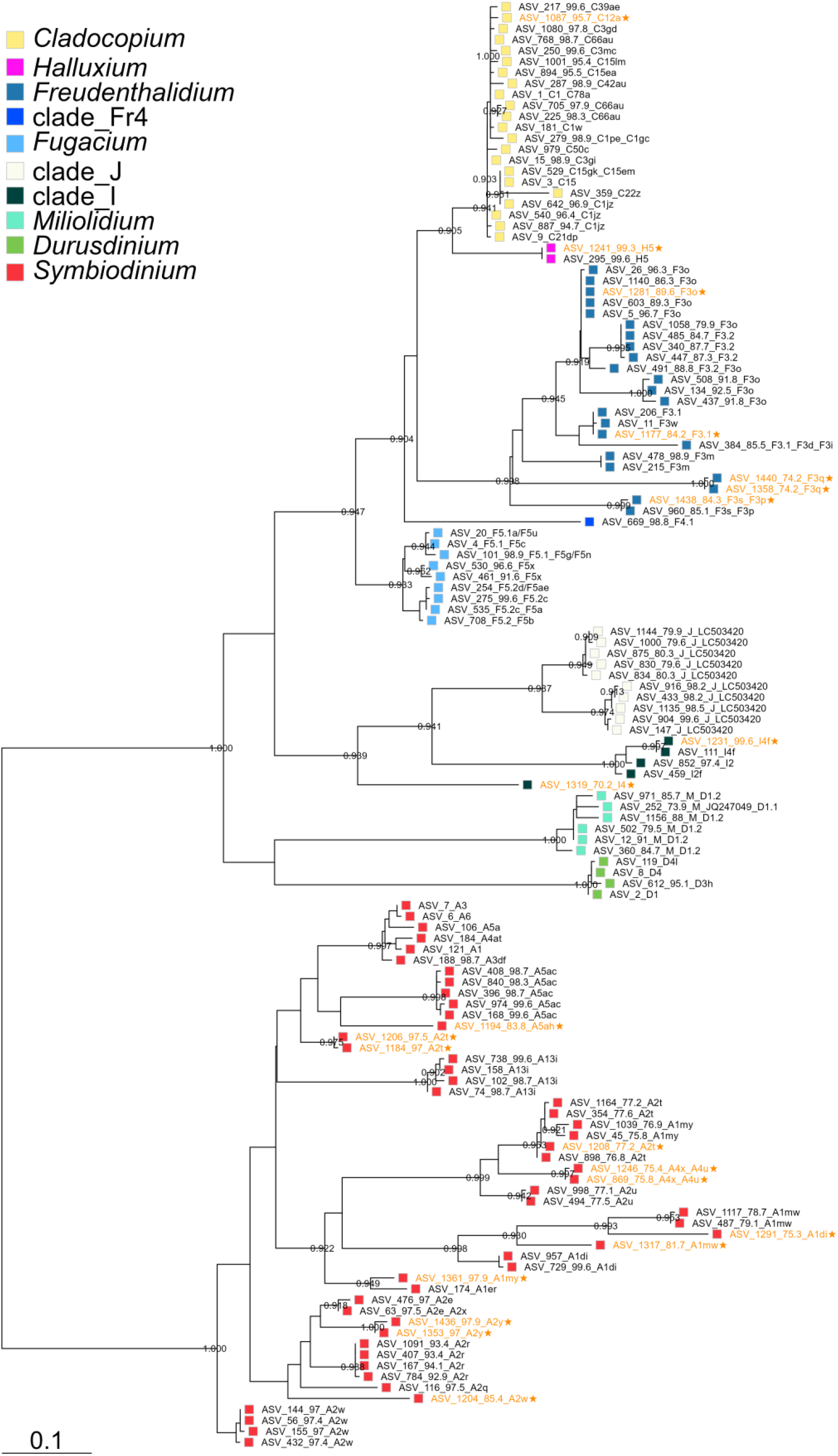
Phylogenetic tree of ASVs used for calculating Faith’s phylogenetic diversity (PD). The ASVs that were filtered out due to abundance are labeled with an asterisk and font in orange. Labels represent ASV numbers and the closest identity match in SymPortal’s database of named DIVs found with vsearch (numbers represent percent identity, not shown if 100%). The tree was built with FastTree from a trimmed alignment (MAFFT + trimAL), unrooted but midpoint-rooted for display, and rooted using *Symbiodinium* as an outgroup for calculating Faith’s PD and PCoA, visualized with the ggtree package. Bootstrap supports were calculated by the default algorithm of FastTree as Shimodaira-Hasegawa (SH)-like local supports, with 1000 resamples, but were not used in calculating weighted UniFrac distances.

**Supplementary Table 3.** Pairwise PERMANOVA results for sample types. Weighted UniFrac distances were calculated in the phyloseq package based on a tree built with FastTree (Fig. S9). P-values were adjusted for multiple testing using the FDR method. *Tridacna* samples were excluded from the DADA2+LULU dataset, and both *Tridacna* and *Amphisorus* samples were excluded from the SymPortal dataset because they violated the assumption of homogeneity of dispersions.

| Comparison | F | R<sup>2</sup> | p | FDR-adjusted p | Significance (FDR) | Pipeline |
| --- | --- | --- | --- | --- | --- | --- |
| Algae_vs_Stichodactyla | 3.637 | 0.342 | 0.095 | 0.114 | - | DADA2 |
| Algae_vs_Porites | 43.624 | 0.799 | 0.001 | 0.00212 | ** | DADA2 |
| Algae_vs_Goniastrea | 39.476 | 0.725 | 0.001 | 0.00212 | ** | DADA2 |
| Algae_vs_Montipora | 3.722 | 0.347 | 0.092 | 0.114 | - | DADA2 |
| Algae_vs_Amphisorus | 19.873 | 0.511 | 0.001 | 0.00212 | ** | DADA2 |
| Algae_vs_Sorites | 5.873 | 0.329 | 0.001 | 0.00212 | ** | DADA2 |
| Algae_vs_Sediment | 0.604 | 0.044 | 0.652 | 0.652 | - | DADA2 |
| Algae_vs_Water | 31.511 | 0.708 | 0.001 | 0.00212 | ** | DADA2 |
| Stichodactyla_vs_Porites | 31.685 | 0.841 | 0.037 | 0.0533 | . | DADA2 |
| Stichodactyla_vs_Goniastrea | 14.904 | 0.598 | 0.056 | 0.0775 | . | DADA2 |
| Stichodactyla_vs_Montipora | 2357.388 | 0.999 | 0.333 | 0.353 | - | DADA2 |
| Stichodactyla_vs_Amphisorus | 1.248 | 0.082 | 0.512 | 0.527 | - | DADA2 |
| Stichodactyla_vs_Sorites | 1.22 | 0.148 | 0.243 | 0.265 | - | DADA2 |
| Stichodactyla_vs_Sediment | 2.275 | 0.221 | 0.093 | 0.114 | - | DADA2 |
| Stichodactyla_vs_Water | 35.556 | 0.816 | 0.026 | 0.0425 | * | DADA2 |
| Porites_vs_Goniastrea | 46.933 | 0.77 | 0.002 | 0.00379 | ** | DADA2 |
| Porites_vs_Montipora | 31.829 | 0.841 | 0.033 | 0.0495 | * | DADA2 |
| Porites_vs_Amphisorus | 122.849 | 0.872 | 0.001 | 0.00212 | ** | DADA2 |
| Porites_vs_Sorites | 35.121 | 0.761 | 0.004 | 0.0072 | ** | DADA2 |
| Porites_vs_Sediment | 33.66 | 0.737 | 0.001 | 0.00212 | ** | DADA2 |
| Porites_vs_Water | 25.914 | 0.683 | 0.002 | 0.00379 | ** | DADA2 |
| Goniastrea_vs_Montipora | 15.001 | 0.6 | 0.032 | 0.0495 | * | DADA2 |
| Goniastrea_vs_Amphisorus | 79.444 | 0.783 | 0.001 | 0.00212 | ** | DADA2 |
| Goniastrea_vs_Sorites | 21.606 | 0.59 | 0.001 | 0.00212 | ** | DADA2 |
| Goniastrea_vs_Sediment | 34.631 | 0.684 | 0.001 | 0.00212 | ** | DADA2 |
| Goniastrea_vs_Water | 46.034 | 0.742 | 0.001 | 0.00212 | ** | DADA2 |
| Montipora_vs_Amphisorus | 1.318 | 0.086 | 0.148 | 0.172 | - | DADA2 |
| Montipora_vs_Sorites | 1.261 | 0.153 | 0.221 | 0.249 | - | DADA2 |
| Montipora_vs_Sediment | 2.326 | 0.225 | 0.081 | 0.108 | - | DADA2 |
| Montipora_vs_Water | 35.78 | 0.817 | 0.019 | 0.0326 | * | DADA2 |
| Amphisorus_vs_Sorites | 8.617 | 0.312 | 0.001 | 0.00212 | ** | DADA2 |
| Amphisorus_vs_Sediment | 16.112 | 0.446 | 0.001 | 0.00212 | ** | DADA2 |
| Amphisorus_vs_Water | 131.402 | 0.868 | 0.001 | 0.00212 | ** | DADA2 |
| Sorites_vs_Sediment | 5.093 | 0.281 | 0.001 | 0.00212 | ** | DADA2 |
| Sorites_vs_Water | 38.189 | 0.746 | 0.001 | 0.00212 | ** | DADA2 |
| Sediment_vs_Water | 24.859 | 0.64 | 0.001 | 0.00212 | ** | DADA2 |
| Goniastrea_vs_Sediment | 56.227 | 0.778 | 0.001 | 0.0028 | ** | SymPortal |
| Goniastrea_vs_Algae | 47.4 | 0.76 | 0.002 | 0.00467 | ** | SymPortal |
| Goniastrea_vs_Sorites | 53.397 | 0.781 | 0.001 | 0.0028 | ** | SymPortal |
| Goniastrea_vs_Water | 22.058 | 0.58 | 0.001 | 0.0028 | ** | SymPortal |
| Goniastrea_vs_Porites | 73.27 | 0.84 | 0.001 | 0.0028 | ** | SymPortal |
| Goniastrea_vs_Montipora | 16.146 | 0.618 | 0.021 | 0.0309 | * | SymPortal |
| Goniastrea_vs_Stichodactyla | 12.447 | 0.555 | 0.018 | 0.0309 | * | SymPortal |
| Sediment_vs_Algae | 0.871 | 0.063 | 0.396 | 0.396 | - | SymPortal |
| Sediment_vs_Sorites | 14.821 | 0.533 | 0.001 | 0.0028 | ** | SymPortal |
| Sediment_vs_Water | 33.437 | 0.705 | 0.002 | 0.00467 | ** | SymPortal |
| Sediment_vs_Porites | 373.691 | 0.969 | 0.001 | 0.0028 | ** | SymPortal |
| Sediment_vs_Montipora | 88.657 | 0.917 | 0.021 | 0.0309 | * | SymPortal |
| Sediment_vs_Stichodactyla | 33.121 | 0.805 | 0.021 | 0.0309 | * | SymPortal |
| Algae_vs_Sorites | 15.36 | 0.561 | 0.003 | 0.006 | ** | SymPortal |
| Algae_vs_Water | 32.844 | 0.716 | 0.001 | 0.0028 | ** | SymPortal |
| Algae_vs_Porites | 438.243 | 0.976 | 0.001 | 0.0028 | ** | SymPortal |
| Algae_vs_Montipora | 110.461 | 0.94 | 0.031 | 0.0347 | * | SymPortal |
| Algae_vs_Stichodactyla | 45.179 | 0.866 | 0.027 | 0.0329 | * | SymPortal |
| Sorites_vs_Water | 63.026 | 0.829 | 0.001 | 0.0028 | ** | SymPortal |
| Sorites_vs_Porites | 1401.653 | 0.992 | 0.003 | 0.006 | ** | SymPortal |
| Sorites_vs_Montipora | 689.549 | 0.99 | 0.018 | 0.0309 | * | SymPortal |
| Sorites_vs_Stichodactyla | 664.601 | 0.99 | 0.026 | 0.0329 | * | SymPortal |
| Water_vs_Porites | 119.051 | 0.908 | 0.001 | 0.0028 | ** | SymPortal |
| Water_vs_Montipora | 24.968 | 0.757 | 0.025 | 0.0329 | * | SymPortal |
| Water_vs_Stichodactyla | 6.776 | 0.459 | 0.023 | 0.0322 | * | SymPortal |
| Porites_vs_Montipora | 14.082 | 0.701 | 0.031 | 0.0347 | * | SymPortal |
| Porites_vs_Stichodactyla | 110.196 | 0.948 | 0.038 | 0.0409 | * | SymPortal |
| Montipora_vs_Stichodactyla | 32.879 | 0.943 | 0.333 | 0.346 | - | SymPortal |

**Supplementary Table 4.** Comparison between alpha diversity measures of different sample types on different datasets (Wilcoxon test).

| Measure | Dataset | Sample type 1 | Sample type 2 | FDR-adjusted *p* | Significance (FDR) | Pipeline |
| --- | --- | --- | --- | --- | --- | --- |
| Chao1 | Filtered | Foraminifera | Environment | 3.70E-07 | *** | DADA2 |
| Chao1 | Filtered | Multicellular hosts | Environment | 1.60E-09 | *** | DADA2 |
| Chao1 | Filtered | Multicellular hosts | Foraminifera | 0.061 | ns | DADA2 |
| Chao1 | Pre_lulu | Foraminifera | Environment | 8.50E-05 | *** | DADA2 |
| Chao1 | Pre_lulu | Multicellular hosts | Environment | 6.30E-04 | *** | DADA2 |
| Chao1 | Pre_lulu | Multicellular hosts | Foraminifera | 0.025 | * | DADA2 |
| Chao1 | Unfiltered | Foraminifera | Environment | 1.40E-07 | *** | DADA2 |
| Chao1 | Unfiltered | Multicellular hosts | Environment | 1.60E-09 | *** | DADA2 |
| Chao1 | Unfiltered | Multicellular hosts | Foraminifera | 0.046 | * | DADA2 |
| FaithPD | Filtered | Foraminifera | Environment | 3.60E-08 | *** | DADA2 |
| FaithPD | Filtered | Multicellular hosts | Environment | 2.40E-09 | *** | DADA2 |
| FaithPD | Filtered | Multicellular hosts | Foraminifera | 0.594 | ns | DADA2 |
| FaithPD | Pre_lulu | Foraminifera | Environment | 2.80E-11 | *** | DADA2 |
| FaithPD | Pre_lulu | Multicellular hosts | Environment | 1.70E-14 | *** | DADA2 |
| FaithPD | Pre_lulu | Multicellular hosts | Foraminifera | 0.612 | ns | DADA2 |
| FaithPD | Unfiltered | Foraminifera | Environment | 4.20E-08 | *** | DADA2 |
| FaithPD | Unfiltered | Multicellular hosts | Environment | 2.40E-09 | *** | DADA2 |
| FaithPD | Unfiltered | Multicellular hosts | Foraminifera | 0.595 | ns | DADA2 |
| Shannon | Filtered | Foraminifera | Environment | 1.20E-06 | *** | DADA2 |
| Shannon | Filtered | Multicellular hosts | Environment | 9.70E-08 | *** | DADA2 |
| Shannon | Filtered | Multicellular hosts | Foraminifera | 0.992 | ns | DADA2 |
| Shannon | Pre_lulu | Foraminifera | Environment | 6.80E-08 | *** | DADA2 |
| Shannon | Pre_lulu | Multicellular hosts | Environment | 3.00E-06 | *** | DADA2 |
| Shannon | Pre_lulu | Multicellular hosts | Foraminifera | 0.002 | ** | DADA2 |
| Shannon | Unfiltered | Foraminifera | Environment | 1.20E-06 | *** | DADA2 |
| Shannon | Unfiltered | Multicellular hosts | Environment | 7.90E-08 | *** | DADA2 |
| Shannon | Unfiltered | Multicellular hosts | Foraminifera | 1 | ns | DADA2 |
| Simpson | Filtered | Foraminifera | Environment | 9.30E-06 | *** | DADA2 |
| Simpson | Filtered | Multicellular hosts | Environment | 1.60E-06 | *** | DADA2 |
| Simpson | Filtered | Multicellular hosts | Foraminifera | 0.851 | ns | DADA2 |
| Simpson | Pre_lulu | Foraminifera | Environment | 7.30E-07 | *** | DADA2 |
| Simpson | Pre_lulu | Multicellular hosts | Environment | 3.00E-06 | *** | DADA2 |
| Simpson | Pre_lulu | Multicellular hosts | Foraminifera | 0.046 | * | DADA2 |
| Simpson | Unfiltered | Foraminifera | Environment | 9.40E-06 | *** | DADA2 |
| Simpson | Unfiltered | Multicellular hosts | Environment | 1.60E-06 | *** | DADA2 |
| Simpson | Unfiltered | Multicellular hosts | Foraminifera | 0.875 | ns | DADA2 |
| Chao1 | Filtered | Foraminifera | Environment | 2.20E-08 | *** | SymPortal |
| Chao1 | Filtered | Multicellular hosts | Environment | 2.50E-09 | *** | SymPortal |
| Chao1 | Filtered | Multicellular hosts | Foraminifera | 0.609 | ns | SymPortal |
| Chao1 | Unfiltered | Foraminifera | Environment | 2.20E-08 | *** | SymPortal |
| Chao1 | Unfiltered | Multicellular hosts | Environment | 2.50E-09 | *** | SymPortal |
| Chao1 | Unfiltered | Multicellular hosts | Foraminifera | 0.694 | ns | SymPortal |
| FaithPD | Filtered | Foraminifera | Environment | 3.40E-08 | *** | SymPortal |
| FaithPD | Filtered | Multicellular hosts | Environment | 1.70E-14 | *** | SymPortal |
| FaithPD | Filtered | Multicellular hosts | Foraminifera | 0.479 | ns | SymPortal |
| FaithPD | Unfiltered | Foraminifera | Environment | 1.50E-12 | *** | SymPortal |
| FaithPD | Unfiltered | Multicellular hosts | Environment | 1.70E-14 | *** | SymPortal |
| FaithPD | Unfiltered | Multicellular hosts | Foraminifera | 0.483 | ns | SymPortal |
| Shannon | Filtered | Foraminifera | Environment | 2.10E-10 | *** | SymPortal |
| Shannon | Filtered | Multicellular hosts | Environment | 6.30E-12 | *** | SymPortal |
| Shannon | Filtered | Multicellular hosts | Foraminifera | 0.049 | * | SymPortal |
| Shannon | Unfiltered | Foraminifera | Environment | 2.10E-10 | *** | SymPortal |
| Shannon | Unfiltered | Multicellular hosts | Environment | 4.60E-12 | *** | SymPortal |
| Shannon | Unfiltered | Multicellular hosts | Foraminifera | 0.049 | * | SymPortal |
| Simpson | Filtered | Foraminifera | Environment | 6.70E-09 | *** | SymPortal |
| Simpson | Filtered | Multicellular hosts | Environment | 1.10E-09 | *** | SymPortal |
| Simpson | Filtered | Multicellular hosts | Foraminifera | 0.112 | ns | SymPortal |
| Simpson | Unfiltered | Foraminifera | Environment | 6.70E-09 | *** | SymPortal |
| Simpson | Unfiltered | Multicellular hosts | Environment | 1.10E-09 | *** | SymPortal |
| Simpson | Unfiltered | Multicellular hosts | Foraminifera | 0.116 | ns | SymPortal |

**Supplementary Table 5.**
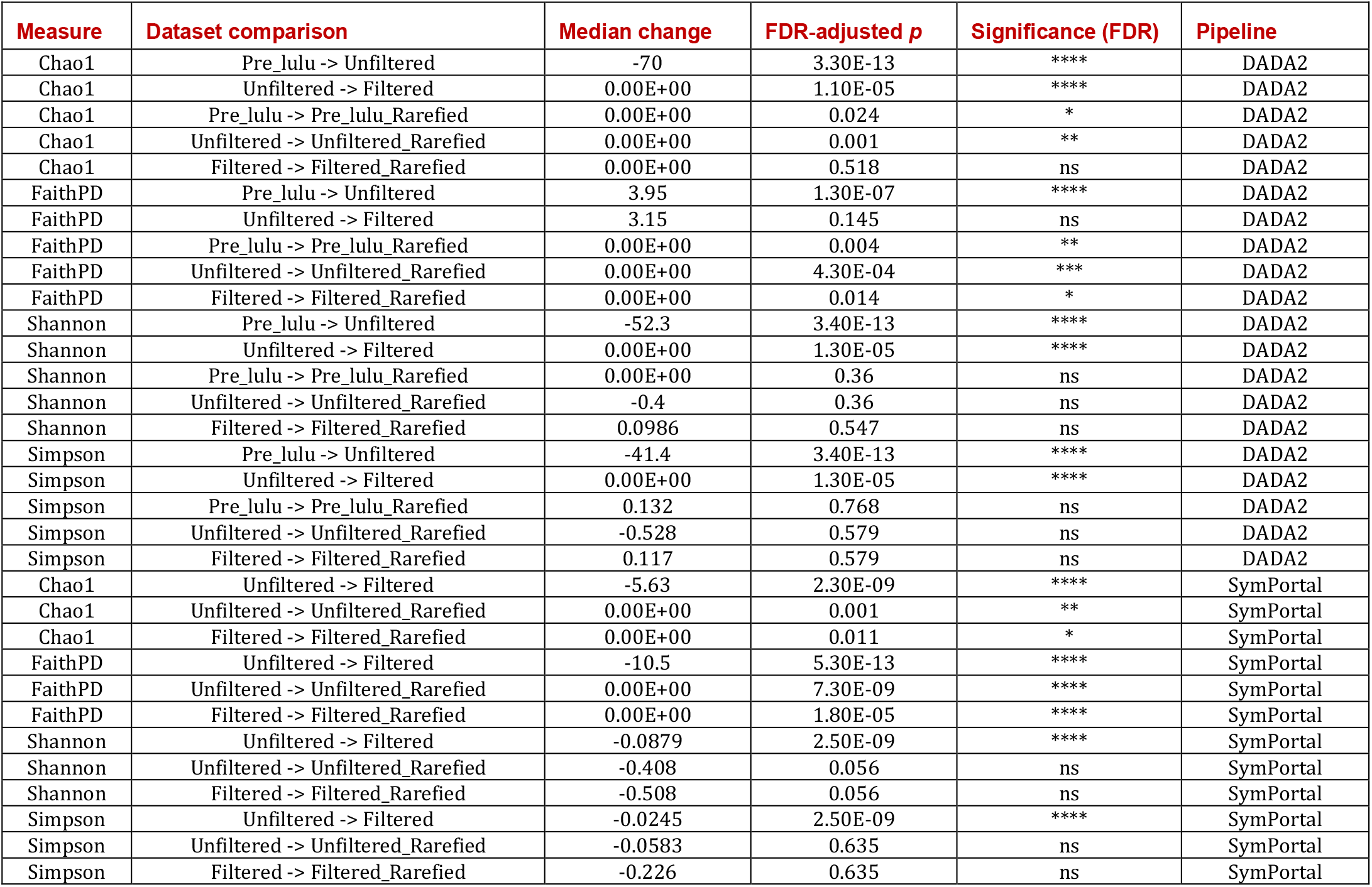
Comparison between diversity measures depending on the filtering and rarefaction (paired Wilcoxon test).

**Supplementary Figure 9.**
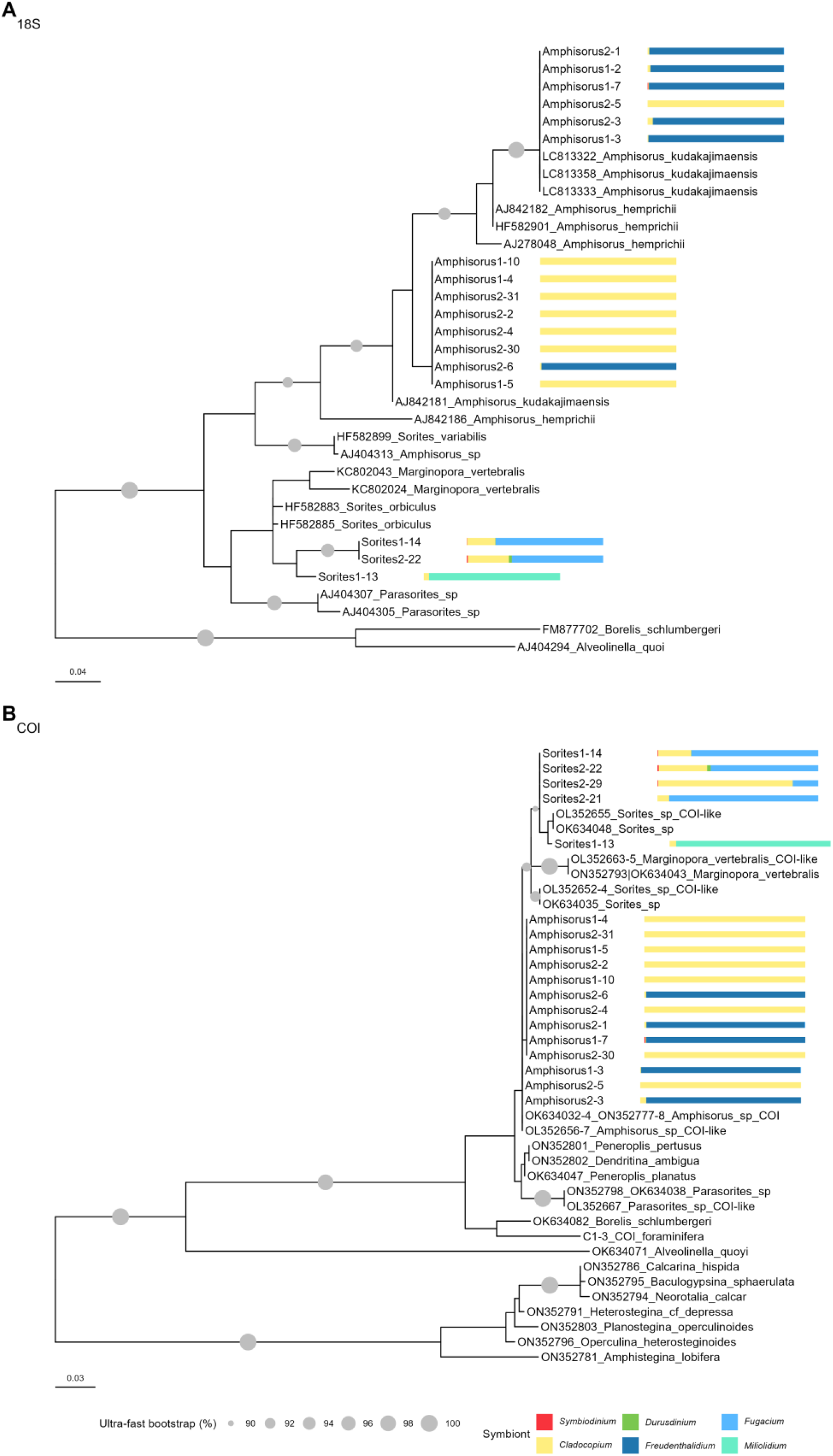
Phylogenetic trees (maximum likelihood) showing the position of foraminiferal samples according to the partial 18S rDNA (A) and COI (B) markers. Only ultrafast bootstrap support values >=90% are shown. Sequences for *Amphisorus* and *Sorites* obtained in this study are shown without GenBank accession numbers and with a bar representing their Symbiodiniaceae composition. Sequence “C3-1_COI_foraminifera” originated from the only *Goniastrea* sample dominated by C1 *Cladocopium* when applying foraminifera-specific COI primers to check if this sample was accidentally swapped with *Amphisorus*. The trees were built as unrooted, but the tree view was modified with the function “Root at mid-point”. Rotaliida were not included in (A) due to the high diversity of the examined 18S region. Both fragments had a length of around 324 bp in our samples. For COI fragment, *Amphisorus* samples differ by one nucleotide only, and this nucleotide was not sequenced in Amphisorus2-1, Amphisorus1-7, Amphisorus2-30.

**Supplementary Table 6.** Accession numbers of marker genes sequenced for the Symbiodiniaceae hosts. The alphanumeric sample code designates the initial sample name upon collection. The sequences labeled with a star are flagged as “unverified” by GenBank and are not included in NCBI BLAST databases. Although these sequences represent PCR product of Miliolida_COI_fwd1 + Miliolida_COI_rev primer pair, similar to some reference sequences such as ON352778.1, the translation of these sequences with the genetic code suggested by the primer authors [47] (transl_table=4) results in the stop codon in the middle of the sequence; whether or not these sequences indeed represent an intact COI gene that is actively transcribed is unclear. Given the missing CDS annotation in GenBank, these sequences are being labeled as “unverified”.

| Sample name | Sample code | GB Accession number | Primer pair |
| --- | --- | --- | --- |
| Montipora2-6 | C2.6 | PV984354 | ZCOI + ZCOIR |
| Montipora1-2 | C1.2 | PV984355 | LC01490 + HC02198 |
| Porites2-1 | C2.1 | PV984519 | ZCOI + ZCOIR |
| Porites1-12 | C1.12 | PV984520 | LC01490 + HC02198 |
| Porites1-6 | C1.6 | PV984521 | LC01490 + HC02198 |
| Tridacna1 | TN1a | PV988022 | 16sar + 16sbr |
| Tridacna2 | TN2a | PV988023 | 16sar + 16sbr |
| Tridacna3 | TN3a | PV988024 | 16sar + 16sbr |
| Tridacna4 | TN4a | PV988025 | 16sar + 16sbr |
| Tridacna5 | TN5a | PV988026 | 16sar + 16sbr |
| Tridacna6 | TN6a | PV988027 | 16sar + 16sbr |
| Tridacna7 | TN7a | PV988028 | 16sar + 16sbr |
| Tridacna8 | TN8a | PV988029 | 16sar + 16sbr |
| Tridacna9 | TN9a | PV988030 | 16sar + 16sbr |
| Amphisorus2-31 | F2.31 | PV988031 | s14F3 + s17 |
| Amphisorus2-31 | F2.30 | PV988032 | s14F3 + s17 |
| Amphisorus2-6 | F2.6 | PV988033 | s14F3 + s17 |
| Amphisorus2-5 | F2.5 | PV988034 | s14F3 + s17 |
| Amphisorus2-4 | F2.4 | PV988035 | s14F3 + s17 |
| Amphisorus2-3 | F2.3 | PV988036 | s14F3 + s17 |
| Amphisorus2-2 | F2.2 | PV988037 | s14F3 + s17 |
| Amphisorus2-1 | F2.1 | PV988038 | s14F3 + s17 |
| Amphisorus1-10 | F1.10 | PV988039 | s14F3 + s17 |
| Amphisorus1-7 | F1.7 | PV988040 | s14F3 + s17 |
| Amphisorus1-5 | F1.5 | PV988041 | s14F3 + s17 |
| Amphisorus1-4 | F1.4 | PV988042 | s14F3 + s17 |
| Amphisorus1-3 | F1.3 | PV988043 | s14F3 + s17 |
| Amphisorus1-2 | F1.2 | PV988044 | s14F3 + s17 |
| Sorites2-22 | F2.22 | PV988045 | s14F3 + s17 |
| Sorites1-14 | F1.14 | PV988046 | s14F3 + s17 |
| Sorites1-13 | F1.13 | PV988047 | s14F3 + s17 |
| Stichodactyla1-1 | An1.1 | PX026258 | ss5+ss3 |
| Stichodactyla1-1 | An1.1 | PX026259 | ITSZ1+ITSZ2 |
| Stichodactyla1-2 | An1.2 | PX026260 | ANEM16SA+ ANEM16SB |
| Amphisorus1-2 | F1.2 | PX092634 * | Miliolida_COI_fwd1 + Miliolida_COI_rev |
| Amphisorus2-31 | F2.31 | PX092635 * | Miliolida_COI_fwd1 + Miliolida_COI_rev |
| Amphisorus2-6 | F2.6 | PX092636 * | Miliolida_COI_fwd1 + Miliolida_COI_rev |
| Amphisorus2-5 | F2.5 | PX092637 * | Miliolida_COI_fwd1 + Miliolida_COI_rev |
| Amphisorus2-3 | F2.3 | PX092638 * | Miliolida_COI_fwd1 + Miliolida_COI_rev |
| Amphisorus2-2 | F2.2 | PX092639 * | Miliolida_COI_fwd1 + Miliolida_COI_rev |
| Amphisorus1-3 | F1.3 | PX092640 * | Miliolida_COI_fwd1 + Miliolida_COI_rev |
| Amphisorus1-4 | F1.4 | PX092641 * | Miliolida_COI_fwd1 + Miliolida_COI_rev |
| Amphisorus1-5 | F1.5 | PX092642 * | Miliolida_COI_fwd1 + Miliolida_COI_rev |
| Amphisorus1-10 | F1.10 | PX092643 * | Miliolida_COI_fwd1 + Miliolida_COI_rev |
| Sorites2-29 | F2.29 | PX092644 * | Miliolida_COI_fwd1 + Miliolida_COI_rev |
| Sorites2-22 | F2.22 | PX092645 * | Miliolida_COI_fwd1 + Miliolida_COI_rev |
| Sorites2-21 | F2.21 | PX092646 * | Miliolida_COI_fwd1 + Miliolida_COI_rev |
| Sorites1-14 | F1.14 | PX092647 * | Miliolida_COI_fwd1 + Miliolida_COI_rev |
| Sorites1-13 | F1.13 | PX092648 * | Miliolida_COI_fwd1 + Miliolida_COI_rev |
| Amphisorus2-1 | F2.1 | PX092649 * | Miliolida_COI_fwd1 + Miliolida_COI_rev |
| Amphisorus1-7 | F1.7 | PX092650 * | Miliolida_COI_fwd1 + Miliolida_COI_rev |
| Amphisorus2-4 | F2.4 | PX092651 * | Miliolida_COI_fwd1 + Miliolida_COI_rev |
| Amphisorus2-30 | F2.30 | PX092652 * | Miliolida_COI_fwd1 + Miliolida_COI_rev |

